# Artificial Hsp100-mediated systems for re-localizing protein aggregates

**DOI:** 10.1101/2022.03.16.484624

**Authors:** Arthur Fischbach, Angela Johns, Xinxin Hao, Kara L. Schneider, Thomas Nyström

**Author notes:** Corresponding authors: T.N., A.F.

## Abstract

Spatial Protein Quality Control (sPQC) sequesters misfolded proteins into specific, organelle-associated inclusions within the cell to harness their toxicity. To approach the role of sPQC in cellular fitness, neurodegenerative diseases and aging, we report on the construction of Hsp100-based systems in yeast cells, which can artificially target protein aggregates to non-canonical locations. We demonstrated that aggregates of mutant Huntingtin (mHtt), the disease-causing agent of Huntington’s disease can be artificially targeted to daughter cells as well as to eisosomes and endosomes with this approach. Removing aggregates from mother cells did not significantly affect their lifespan and targeting mHtt to multiple smaller aggregates rather than one large inclusion did not alter its toxicity. We demonstrated that this approach is able to manipulate mHtt inclusion formation also in human cells and has the potential to be a useful complementation to present therapeutic approaches aimed at alleviating age-related neurodegenerative diseases.

## Introduction

Aging is a ubiquitous process in life affecting most organisms, in particular animals but also many unicellular organisms. It is widely assumed that aging is caused by an accumulation of cellular damage over time (*1-3*). In the budding yeast *Saccharomyces cerevisiae*, the cellular damages considered to contribute to aging include protein aggregates, oxidatively damaged proteins, defective mitochondria and vacuoles, and extra-chromosomal ribosomal DNA circles (ERCs) (*4, 5*). Yeast cells have developed strategies to asymmetrically segregate such damages and toxic molecules during cell division, so that daughter cells (also known as buds) produced during cytokinesis are mostly free from age-related damages. For example, asymmetric inheritance has been shown for damaged and aggregated proteins (*6, 7*), defective mitochondria (*8, 9*), vacuoles (*10*), and ERCs (*11*). While these studies demonstrate an association between the segregation of specific cellular damages and aging/rejuvenation, causal links between such damages and aging have not been firmly established. For example, while genetic manipulations causing an increased accumulation of protein aggregates shortens yeast replicative lifespan and manipulation causing a decreased accumulation of aggregates prolongs lifespan (*12-17*), it is not clear if protein aggregates themselves are the cause of aging as such manipulations are expected to alter a multitude of processes in the cell.

Protein aggregates are formed by a variety of different routes when misfolded proteins; proteins that have lost their tertiary structure, overwhelm the protein quality control (PQC) systems of the cell. Such aggregation is harmful to the cell due to the loss of function of the proteins aggregating and also due to gain of toxic, non-canonical, functions of the aggregates formed. A failure of PQC leads for instance to neurodegenerative diseases in humans, where aggregated proteins impair cellular functions and eventually cause cell death (*18*). In some organisms, protein aggregates can be cleared from the cell by ‘disaggregases’ of the Hsp100 family, such as Hsp104 in yeast (*19*). If such clearance fails, aggregates are sequestered into large inclusions at certain cellular sites in the cell. It has been suggested that this sequestration of misfolded and aggregated proteins into large inclusions at specific sites might decrease the toxicity of such aggregates. This may be due to, for example, a reduction of the exposed surface area restricting interactions with other functional proteins and limiting titration of PQC components, including chaperones (*20, 21*). In addition, it has been shown that inclusions associated with the mitochondria are cleared out faster than inclusions that are not (*22*), indicating that specific locations in the cell may be more efficient in certain PQC processes.

In yeast, misfolded and aggregated proteins, when formed, first accumulate at multiple sites, known as stress foci, CytoQs or Q-bodies throughout the cytosol and at the surface of various organelles, such as the endoplasmic reticulum (ER), vacuole and mitochondria. Upon prolonged proteostatic stress, aggregates in stress foci coalesce into larger inclusions by an energy-dependent process (*23*). These large inclusions are categorized by their proximity to specific organelles (juxtanuclear quality control site (JUNQ) on the cytosolic side of the nuclear membrane (*24*); intranuclear quality control site (INQ) in close proximity to the JUNQ, but within the nucleus, next to the nucleolus (*25*); peripheral vacuole-associated insoluble protein deposit (IPOD) proximal to the vacuole (*23, 24*); and a site adjacent to mitochondria (*26*)). Different sorting mechanisms and factors appear to play a role in the sorting of misfolded proteins to each specific site (*25, 27, 28*) but it appears that most misfolded proteins studied are sequestered to all these sites (*23, 24, 29*). However, the vacuole-associated IPOD (*30*) seems to be the deposition site for amyloid proteins including the protein mutant Huntingtin, which is the causative agent of Huntington’s disease (HD) (*24*). In contrast, mammalian cells sequester aggregated proteins in a deposition site close to the centrosome, known as the aggresome (*31, 32*).

That spatial control of aggregates is important for cellular fitness, rejuvenation, and aging is inferred from results using mutations causing defects in spatial PQC (sPQC). This study was aimed at generating a different approach to studying the importance of sPQC by artificially transporting protein aggregates to non-canonical, but targeted, locations in the cell by using protein fusions to the yeast aggregate-binding protein Hsp104. We demonstrate that this approach is successful in forming inclusions at non-canonical sites in both yeast and human cells and has the potential to be useful as an alternative, complementary strategy to study the role of sPQC in aging and neurodegenerative disease.

## Results and Discussion

### Generation of an artificial aggregate targeting system (ATS) to transport aggregates into daughter cells

Hsp104 has been shown to recognize stress-induced and age-related protein aggregates and to contribute to their retention in the mother cell (*7, 33, 34*). However, the role of this mother cell retention of aggregates is not clear but has been suggested to contribute to the aging of mother cells and the rejuvenation of daughters (*6, 7, 35, 36*). In order to further elucidate the role of such retention of aggregates, and their sequestration to certain organelle-associated sites in the cell (i.e. IPOD, JUNQ, INQ and mitochondria) (*23-26*), we aimed to remove aggregates from mother cells during cytokinesis by targeting them to daughter cells. In order to do so, we first fused Hsp104-GFP to the myosin motor protein Myo2 arguing that Hsp104 fused to Myo2 could bring aggregates to daughter cells by anterograde transport along actin cables. While this system has shown successful targeting to the bud, it led in many cases to a lethal two-budded phenotype (fig. S1A). In a second attempt, we instead fused Hsp104-GFP to Pea2, a protein component that is transported with Myo2 along actin cables to the polarisome protein complex at the tip of the bud (*37*). The *HSP104-GFP-PEA2* fusion gene was integrated genomically into the *MET15* locus, leaving the endogenous *HSP104* and *PEA2* genes unaltered (Fig. 1A). This approach showed successful targeting to the bud (Fig. 1B) and we refer to this Hsp104-Pea2 chimera system as ‘Aggregate Targeting System 1’ (ATS1; Fig. 1, A and C). Interestingly, Hsp104-GFP-Pea2 forms a single inclusion, indicating that most Hsp104-GFP-Pea2 is sequestered into the same site, suggesting that the specific targeting of Hsp104 may cause molecular crowding/seeding that induces Hsp104 inclusion formation. This is different from the normal, mainly cytosolic, localization of Hsp104 (Fig. 1B). To test the hypothesis of a seed/crowding-triggered inclusion formation, we induced artificial crowding of Hsp104 by fusing it to the p53 tetramerization domain. This triggered in support of our hypothesis the formation of several Hsp104 inclusions per cell in the absence of proteostatic stress (Fig. 1D).

**Fig. 1.**
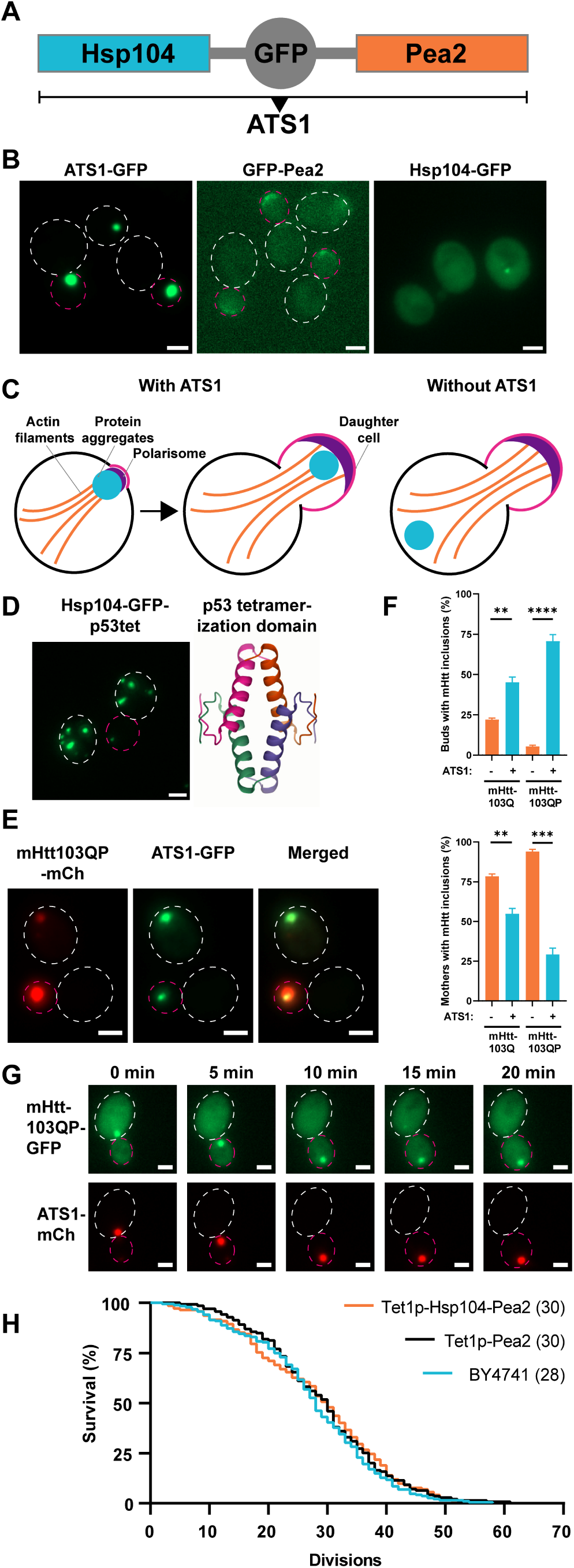
Generation of an aggregate targeting system to the bud. (**A**) Domain composition of aggregate targeting system 1 (ATS1). (**B**) Representative fluorescence microscopy images of ATS1 localizing to the daughter cell (Daughters are indicated by a dashed circle in magenta. Mothers are indicated by a dashed circle in white). Scale bar represents 2 µm. (**C**) Conceptual schematic of ATS1. (**D**) Left panel: Fusing the p53 tetramerization domain (p53tet) to Hsp104 leads to inclusion formation. Hsp104-GFP-p53tet was under the control of an *ADH1* promoter. Right panel: Crystal structure of four p53tet molecules assembled to a tetramer (PDB: 1AIE). (**E**) mHtt103QP aggregate transport into buds after 4.5 h induction with galactose in presence of ATS1. (**F**) Quantification of buds or mothers with mHtt inclusions from (E). (**G**) Time lapse microscopy images of mHtt103QP-GFP aggregates (constitutive expression from GPD promoter) entering the bud using ATS1-mCherry. (**H**) Replicative lifespan analysis via micromanipulation of cells with inducible ATS1. Median lifespan is indicated in parentheses.

To investigate if ATS1 can recognize and transport protein aggregates into the bud, we expressed the misfolding proteins mutant Huntingtin 103Q and 103QP (mHtt103Q and mHtt103QP, respectively). mHtt103QP is the Huntington’s disease associated N-terminal fragment of human huntingtin and exhibits an expansion from a 25 to a 97 poly-glutamine stretch leading to misfolding and aggregation (*38*). mHtt103Q lacks the adjacent proline-rich region. We found that ATS1 could recognize mHtt103QP and mHtt103Q aggregates, both when the gene was under control of an inducible *GAL1*-promoter (Fig. 1E) or a constitutive promoter (fig. S1B), with up to 71% of daughters containing mHtt inclusions and an up to 65% reduction of mother cells containing mHtt inclusions (Fig. 1F). A time-lapse example of ATS1-dependent binding and transport of mHtt103QP-GFP aggregates into the bud is shown in Fig. 1G and movie S1. We term this general principle of targeting proteins to an artificial inclusion via chimeras ‘INTACs’ (inclusion-targeting chimeras), in analogy to proteolysis-targeting chimeras (PROTACs) (*39, 40*) or lysosome-targeting chimeras (LYTACs) (*41*).

In addition to mHtt, we investigated if ATS1 can bind and transport other misfolding model proteins using the temperature-sensitive alleles *guk1-7, pro3-1* and *gus1-3* (*42*). All these misfolding model proteins showed inclusions co-localizing with ATS1 and were transported into the bud, indicating a general ability of ATS1 to bind misfolded proteins forming aggregates (fig. S1, C to E).

To check if ATS1 caused cellular defects, we measured growth using a drop test and the activity of a reporter of the Hsf1-dependent stress response (*43*). We could not detect any growth defects for the ATS1 strain (fig. S1F) and only a slight increase in cellular Hsf1 activity in comparison to controls (fig. S1G).

To investigate if the artificial ATS1 inclusions are functional, we tested whether aggregates in the inclusions were cleared with a similar rate as aggregates in wild type cells using the misfolding protein pro3-1. We found that pro3-1 aggregates were cleared with a similar rate as aggregates in cells without ATS1, demonstrating successful disaggregation as well as functionality of Hsp104 in the ATS1 inclusions (fig. S1, H and I).

With the development of the ATS1 tool of aggregate removal from the mother cell, we had the unique opportunity to test if protein inclusions might constitute a causal factor in the replicative aging process of yeast. We performed microdissection lifespan analysis with the constitutive expression of ATS1 and could observe a small but not statistically significant effect (p = 0.38) on the lifespan of ATS1 mother cells compared to mother cells without ATS1 (fig. S1J). Similar results were obtained with microfluidics-based lifespan experiments (fig. S1K). Because in constitutive ATS1 strains, the daughters picked for lifespan analysis are born with more damage than they would otherwise have, we used a doxycycline-inducible ATS1 to overcome this problem so that the transport system was induced after picking the daughters for lifespan analysis. However, this approach gave similar results as with the constitutive ATS1 that was not statistically significant (p = 0.83) (Fig. 1H).

There are several possibilities explaining the small or non-existent effect of mother cell lifespan when freed of aggregates. Firstly, the large, single inclusions formed by the artificial ATS1 might be mostly inert and non-toxic whereas smaller, and potentially harmful, oligomers and aggregates of misfolded proteins not recognized by ATS1 still stay in the mother cell. Secondly, it is possible that the efficiency of ATS1 in transporting aggregates might diminish with age as it relies on actin cables, which have been shown to degenerate during mother cell aging (*44*).

### Hsp70s as well as Hsp104 oligomerization and ATP binding are essential for ATS1 formation

To investigate which genes are important for ATS1 formation, we performed single-deletions of several proteostasis genes, but none showed an effect on inclusion formation (fig. S2A). Additionally, deletions of diffusion barrier genes did not markedly change inclusion formation or targeting of ATS1 to the bud (fig. S2B). The only genetic perturbation that markedly affected ATS1 was a double deletion of the cytosolic Hsp70 genes, *SSA1* and *SSA2*, which highly reduced ATS1 formation (Fig. 2A).

**Fig. 2.**
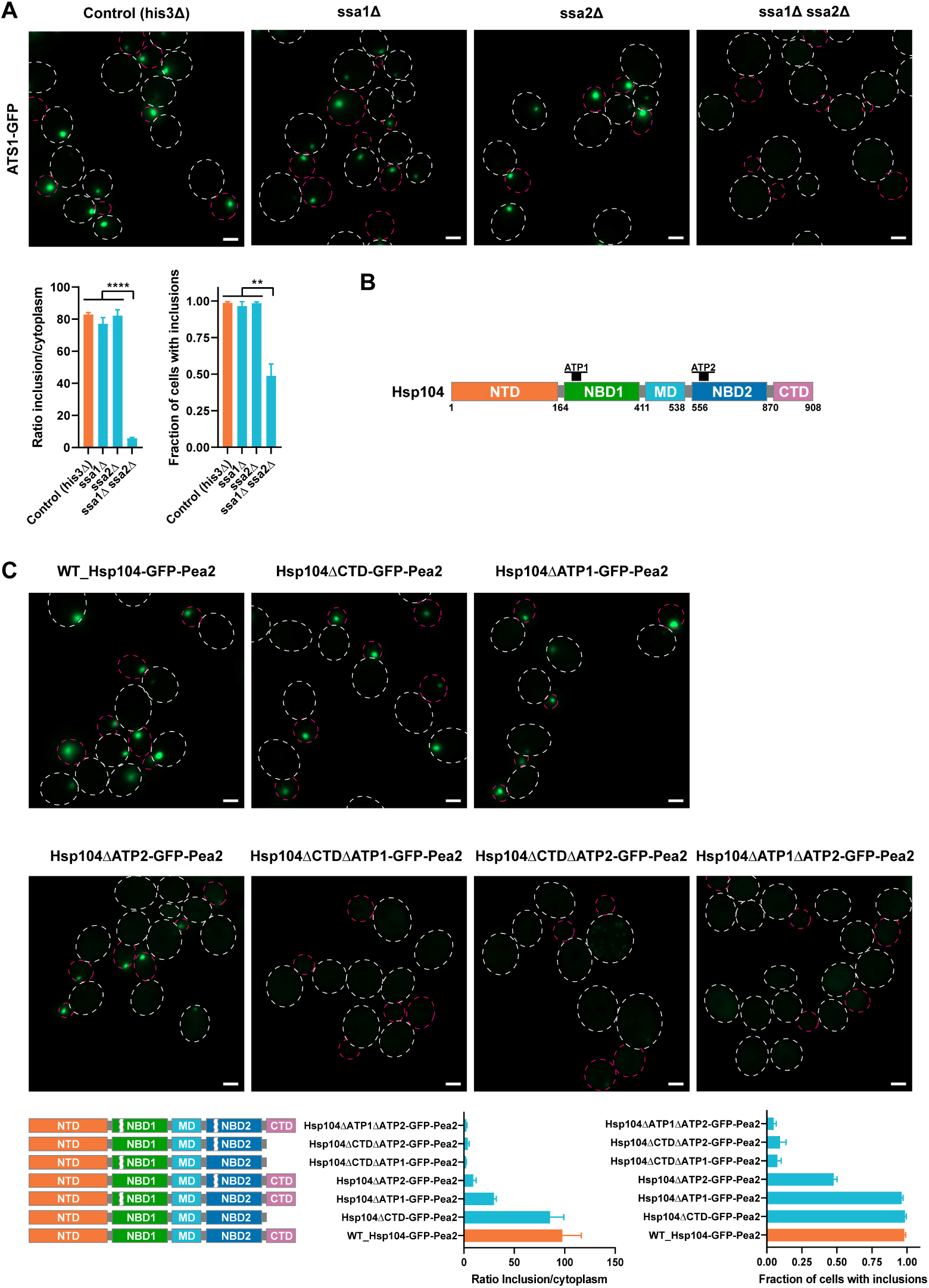
Genes and genetic modifications affecting ATS1. (**A**) Top panel: Representative fluorescence microscopy images showing the effect of double deletion of the Hsp70 genes *SSA1* and *SSA2* on ATS1-GFP formation. Cells were grown at 22 °C. Bottom panel: Quantifications of the GFP intensity ratio of inclusion to cytoplasm. Quantifications of the fraction of cells with ATS1 inclusions. (**B**) Domain architecture of Hsp104. Hsp104 is composed of an N-terminal domain (NTD), nucleotide-binding domain 1 (NBD1), middle domain (MD), nucleotide-binding domain 2 (NBD2), and C-terminal domain (CTD). NBD1 and NBD2 harbour the ATP binding sites ATP1 and ATP2, respectively. (**C**) Top panel: Representative fluorescence microscopy images showing the influence of the NBD and oligomerization of Hsp104 on inclusion formation of ATS1. Cells were grown at 30 °C. Bottom panel: Quantifications of the GFP intensity ratio of inclusion to cytoplasm. Quantifications of the fraction of cells with inclusions.

Further, we investigated which regions in Hsp104 were important for artificial inclusion formation of ATS1. Hsp104 assembles into a homohexamer, forming a ring with a central pore. Thus, we assumed that oligomerization domains of Hsp104 are important for inclusion formation. The C-terminal domain (CTD) has been shown to play a role in oligomer assembly (*45*). However, deletion of the CTD showed no effect on ATS formation (Fig. 2, B and C). A previous study investigated *in vitro* the role of the nucleotide binding domains NBD1 and NBD2 in oligomer assembly, with NBD2 showing an influence on Hsp104 oligomerization whereas the NBD1 did not (*46*). We deleted the ATP binding sites ATP1 or ATP2 (8 amino acid deletions each) in both NBDs and checked for ATS1 formation. In accordance with Schirmer et al. (*46*), the ATP2 deletion showed a high reduction of ATS1 formation whereas the ATP1 deletion showed a less pronounced reduction (Fig. 2C). This indicates that ATP1, in addition to ATP2, plays an *in vivo* role in Hsp104 oligomerization. If the ATP binding site deletions were combined with the deletion of the CTD, ATS1 formation was further decreased, demonstrating a supportive role of the CTD for Hsp104 oligomerization. The strongest reduction in ATS1 formation was achieved by deleting ATP1 and ATP2 together, indicating that ATPase function and catalytic activity of Hsp104 are closely tied to inclusion formation (Fig. 2C). Taken together, these results demonstrate that the presence of Hsp70 chaperones as well as the oligomerization and ATP binding ability of Hsp104 are essential for ATS1 formation.

### ATS1 sequesters endogenous Hsp104 and is enriched for various proteostasis factors

To characterize ATS1 further, we checked for enrichment/co-localization of proteostasis factors. First we tested for co-localization with endogenous Hsp104. Interestingly, ATS1 sequestered most of the endogenous Hsp104 in the artificial inclusion (Fig. 3A), which explains how all the misfolded model proteins tested accumulate specifically at these sites. This implies that ATS1 incorporates the regular Hsp104 inclusions, e.g. also the ones arising during aging, which have been previously described as age-dependent protein deposits (APOD) (*14, 47*). We confirmed this in old cells using microfluidics (fig. S3A). Furthermore, ATS1 was enriched with other proteostasis factors, such as endogenous Hsp42, a small heat shock protein; Tsa1, a thioredoxin peroxidase; and Ssa1, an Hsp70 protein (Fig. 3, B to D), but not with GFP (Fig. 3E). Also, ATS1 was enriched with the J-proteins Ydj1 and Sis1; with Hsc82, an Hsp90 protein, but not with the ATPase component Sse1. No enrichment with the autophagosome protein Atg8 could be detected (fig. S3, B to F). Some proteins, like Btn2 and Mca1, only co-localized with ATS1 after a heat shock (fig. S3, G and H), i.e. when aggregates accumulate at ATS1.

**Fig. 3.**
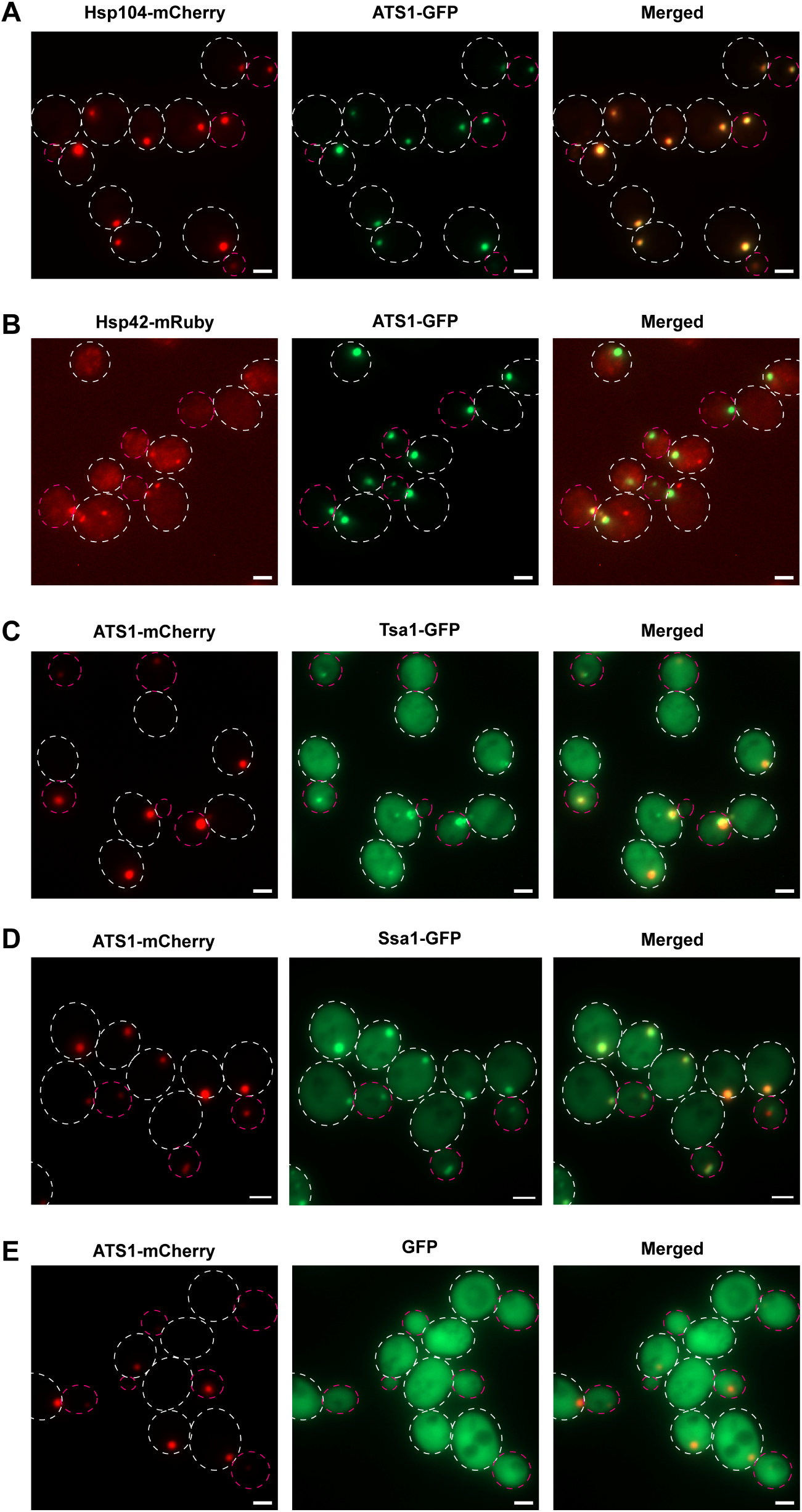
Endogenous proteins co-localizing with ATS1. (**A**) Representative fluorescence microscopy images of the co-localization of ATS1 (green, GFP-labeled) with endogenous Hsp104 (red; mCherry-labeled). (**B**) Co-localization of ATS1 (green; GFP-labeled) with endogenous Hsp42 (red; mRuby-labeled). (**C**) Co-localization of ATS1 (red; mCherry-labeled) with endogenous Tsa1 (green; GFP-labeled). (**D**) Co-localization of ATS1 (red; mCherry-labeled) with endogenous Ssa1 (green; GFP-labeled). (**E**): No enrichment of GFP (green; expression from GPD promoter) in ATS1 inclusions (red; mCherry-labeled).

### Artificial sequestration of aggregates to non-canonical sites by other ATS constructions

Since the results above suggested that a seed structure like the polarisome, i.e. molecular crowding, appears sufficient to induce inclusion formation we tried to use this fact to target inclusions to other artificial locations. The endosome forms a seed-like structure and we therefore fused Hsp104 to the endosomal protein Snf7 (ATS2) and found that this led to formation of artificial inclusions (Fig. 4A). In another approach, we fused the eisosome protein Pil1 to Hsp104 (ATS3). The eisosomes form multiple seed-like structures nearby the cell membrane and fusing Pil1 to Hsp104 induced the formation of artificial inclusions (Fig. 4B). Both the artificial ATS2 and ATS3 inclusions were able to sequester mHtt aggregates (Fig. 4, C and D and fig. S4, A and B). As seen in Fig. 4D, the Pil1-Hsp104 fusion resulted in mHtt forming multiple aggregates (‘class III phenotype’: 3 or more aggregate inclusions per cell (*22, 48*)) instead of the typical one to two in wild type cells. We used this system to elucidate the long-standing question if multiple aggregates are more deleterious to the cell than single (few) aggregates due to exposing an increased surface area that may, for example, sequester and deplete chaperones from the cytosol (*20, 21, 49, 50*). However, when determining the growth rate of cells expressing mHtt constitutively (Fig. 4E) or after induction (Fig. 4F) we found no significant effect of mHtt forming multiple aggregates instead of one to two.

**Fig. 4.**
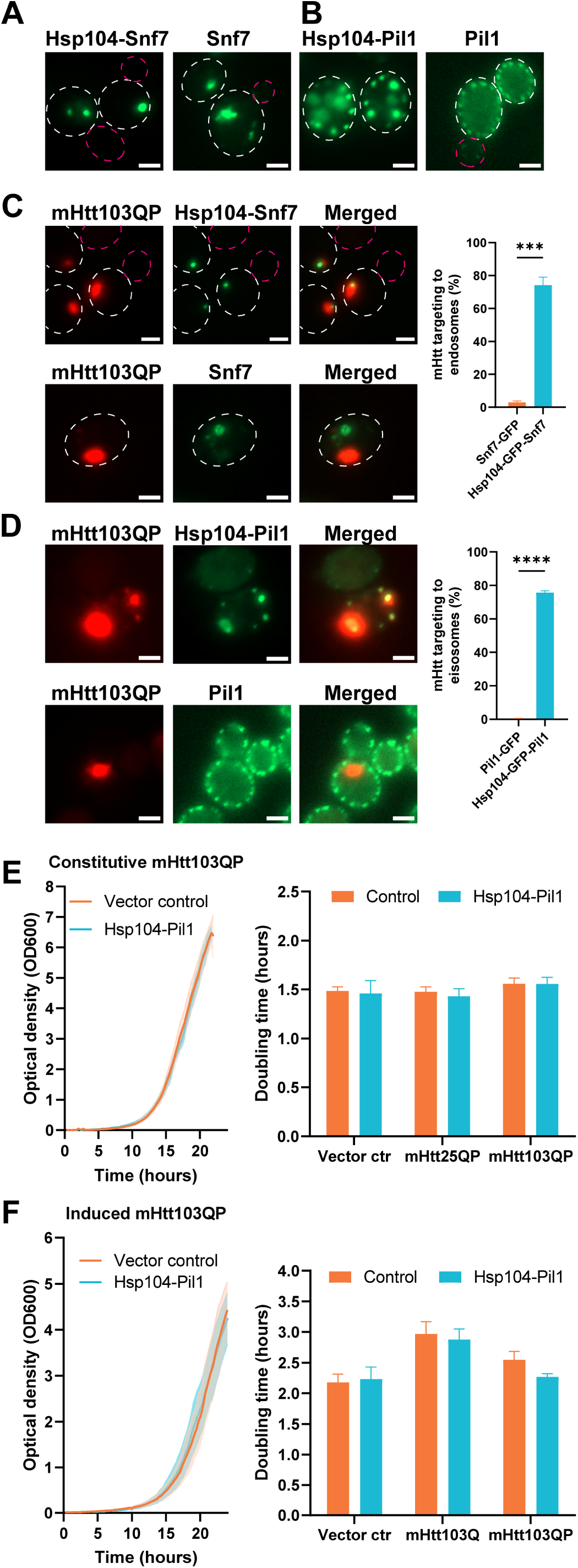
Artificial sequestration of aggregates to non-canonical sites can be achieved by other ATS constructions. (**A**) Representative fluorescence microscopy images showing the targeting of Hsp104 inclusions to the endosome by fusing Hsp104-GFP to Snf7 (ATS2). (**B**) Targeting of Hsp104 inclusions to eisosomes by fusing Hsp104-GFP to Pil1 (ATS2). (**C**) Left panel: Representative fluorescence microscopy images of the artificial targeting of mHtt103QP-mCherry aggregates to endosomes by using Hsp104-GFP-Snf7. Right panel: Quantifications of mHtt targeting efficiency. n = 3 independent experiments. The expression of mHtt103QP-mCherry was induced by galactose for 4.5 h. (**D**) Left panel: Representative fluorescence microscopy images of the artificial targeting of mHtt103QP-mCherry aggregates to eisosomes by using Hsp104-GFP-Pil1. Right panel: Quantifications of mHtt targeting efficiency. n = 3 independent experiments. The expression of mHtt103QP-mCherry was induced by galactose for 4.5 h. (**E** and **F**) Growth curves (left panels) and doubling time (right panels) of cells with artificial targeting of mHtt103QP-mCherry aggregates to the eisosomes by using Hsp104-Pil1. mHtt103QP was under the control of a constitutive GPD promoter (E) or an inducible *GAL1* promoter (F). n = 3-6 independent experiments.

### Screening for ‘inclusion generators’ other than Hsp104

To find proteins other than Hsp104 that can initiate and form artificial inclusions we performed a genome-wide screen using synthetic genetic array (SGA) technology. We devised a novel approach to generate chimeras in a large-scale fashion by fusing Pea2 to a GFP-binding nanobody (GFP binding protein, GBP) (*51*). This construct was crossed into the Yeast-GFP collection (*52*). In that way, 4159 different chimeras were generated, consisting of an endogenous protein bound to Pea2 (Fig. 5A). We identified 71 proteins that can initiate and form artificial inclusions in the cytoplasmic fraction (Fig. 5, B to C and table S1) with false discovery rate (FDR) of < 0.05. We termed this class of proteins ‘inclusion generators’ (IGs) because they have the ability to generate inclusions when a seed is provided (in this case by Pea2). We tested if any of those chimeras were able to transport mHtt103QP into the daughter cell but found that Hsp104 was the only protein capable of doing this (table S1). This suggests that inclusion formation of any protein is not sufficient to sequester the aggregation-prone mHtt to such inclusions and that the mHtt aggregates found to bind the Pea2-Hsp104 fusion are *bona fide* aggregates.

**Fig. 5.**
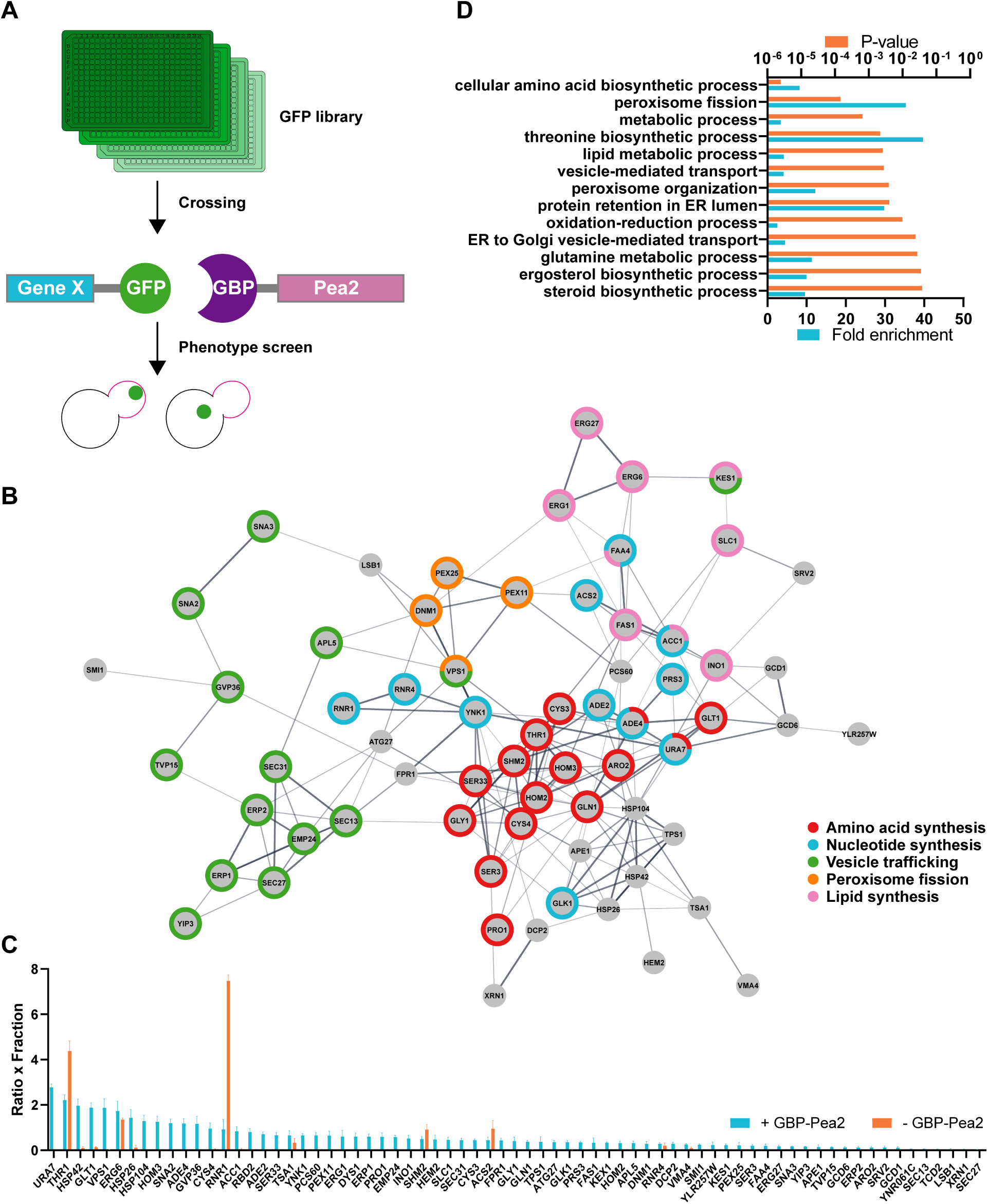
Screening for ‘inclusion generators’ other than Hsp104. (**A**) Scheme for screen strategy. GFP binding protein (GBP) was fused to Pea2 and crossed with the GFP library using SGA technology. Cells were screened for a phenotype with a strong artificial inclusion of a chimera consisting of the GFP-labelled gene of interest and Pea2. (**B**) Functional clustering of hits for inclusion generators. Edges (lines) represent published physical protein–protein interactions and functional associations. Nodes represent the screen hits. (**C**) Screen data output for product of inclusion-to-cytoplasm-ratio and fraction of cells with inclusions (Ratio x Fraction). n = 3 independent experiments. (**D**) Gene ontology biological process analysis of screen hits.

Gene ontology analysis revealed a marked enrichment of genes involved in vesicle trafficking as well as in the synthesis of amino acids, nucleotides and lipids. Interestingly, several peroxisomal, lipid droplet genes, and Dcp2 (a P-body protein) were identified in the screen as inclusion generators, suggesting that their organization and formation might be mediated by a similar seed mechanism (Fig. 5, B and D). Of note, the artificial inclusions of those genes were also transported into daughter cells, suggesting that it is possible to manipulate their spatial location in the cell (fig. S5A). This may be instructive for example in the role of the importance of spatial control in P-body formation.

Interestingly, 33.8% of the here identified IGs were proteins found in a previous screen (*53*) for heat-shock-induced aggregates (p < 0.0001, chi-square test; 6.4 fold enrichment, fig. S5B). In addition, the IGs were enriched for proteins that form homo oligomers (p < 0.0001, chi-square test, 2.7 fold enrichment, table S2), according to calculations based on the Uniprot database and a recent genome-wide oligomerization estimation (*54*). Hetero-oligomerization is probably also a driving force for the IGs, but to our knowledge, no comparative study has been performed.

Many IGs are metabolic and lipid enzymes. It was previously demonstrated that numerous of those can form inclusions or filaments in human cells (*55, 56*) and upon starvation (*57, 58*) and exponential growth of yeast cells (*59, 60*). The physiological role of these inclusions or filaments is unclear, but ATP depletion plays a possible role in their formation (*61*).

We discovered that Tsa1 and the small heat shock proteins Hsp42 and Hsp26 are IGs. Hsp42 and Hsp26, as well as Btn2, have previously been termed ‘sequestrases’ for their ability to sequester misfolded/aggregated proteins (*62, 63*). We did not identify Btn2 as an IG in our screen (rank 395), but this might be due to low expression levels in the absence of stress.

The current view is that Hsp104, Tsa1 or Hsp42 accumulate into cytoplasmic foci upon cellular damage events (*16, 64*), but our results demonstrate that those proteins can also organize and generate self-inclusions in the absence of damage when concentrated to a specific structure/location in the cell. However, Tsa1 and the small heat shock proteins were all unable to sequester mHtt aggregates on their own (in the absence of Hsp104 inclusions; table S1), demonstrating that the formation of self-inclusions is not sufficient to sequester aggregates.

### Manipulations of mHtt inclusions in human cells

To investigate if mHtt inclusions can also be manipulated in cultured human cells, we transfected HEK293 cells with codon-optimized Hsp104. In analogy to our yeast-based engineering, we aimed to induce artificial Hsp104 inclusions in human cells by fusing it to various potential seed-forming proteins. Fusing Hsp104-mCherry to the HIV capsid protein Gag (ATS4) induced the formation of artificial inclusions, while Hsp104-mCherry was distributed diffusely in the cytoplasm (Fig. 6A). Gag is the main structural protein of retroviruses and its expression alone is sufficient to drive the formation of virus-like particles (*65*). Of note, Gag-based virus-like particles have the ability to transport cargo through the plasma membrane (*66*) and potentially protein aggregates. We observed that Hsp104-Gag inclusions could serve as aggregation seeds for mHtt119Q, suggesting that spatial manipulations of disease aggregates are possible with this approach (Fig. 6B). In human cells, the mHtt119Q aggregates are normally localized in the aggresome proximal to both the centrosome and the nucleus (*31, 32*). Spatial manipulations of the aggresome should therefore change its distance to the nucleus. Using Hsp104-Gag for aggregate seeding indeed changed the distance of the early mHtt119Q seed, as determined by timelapse microscopy, showing that mHtt119Q aggregates were diverted from the aggresome (Fig. 6C). In 66 % of the cases, an artificial Hsp104-Gag inclusion served as an aggregation seed for mHtt119Q (Fig. 6D). Using Hsp104-Gag also changed the morphology of mHtt inclusions: There was a higher frequency of several mHtt inclusions and of ‘ruffled’ inclusions (Fig. 6, E and F), further indicating that normal aggresome formation of mHtt119Q had been debilitated. Thus, the system can be used to test to what extent the toxicity of different disease proteins is affected, or not, by being sequestered to aggresomes.

**Fig. 6.**
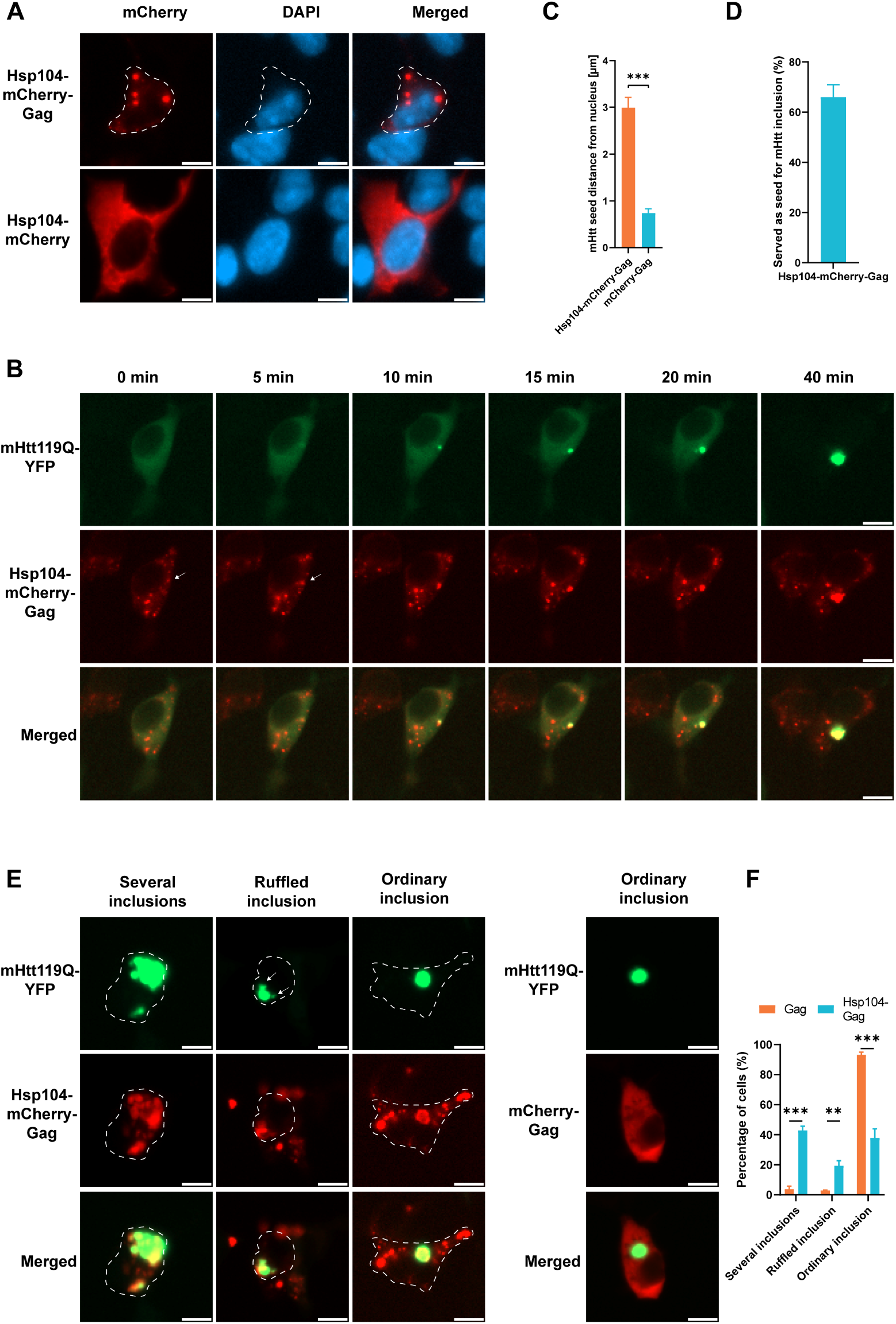
Manipulations of mHtt inclusions in human cells. (**A**) Representative fluorescence microscopy images showing a comparison between HEK293 cells transfected with Hsp104-mCherry or Hsp104-mCherry-Gag. (**B**) Representative fluorescence microscopy images of artificial Hsp104-mCherry-Gag inclusions serving as a seed for mHtt119Q aggregation (marked by a white arrow). (**C**) Quantification of the distance to the nucleus of mHtt aggregate seeds from B. (**D**) Frequency quantification of artificial Hsp104-mCherry-Gag inclusions serving as seed for mHtt119Q aggregation. (**E**) Representative fluorescence microscopy images of inclusion manipulations by Hsp104-mCherry-Gag. Arrows indicate ruffled edges of inclusions. Scale bars represent 10 µm. (**F**) Quantification of E.

Potentially, improved future versions of an ATS can be used as a therapy in patients with Huntington’s disease (HD) or other proteinopathies, as an alternative to current therapy strategies, like antisense oligonucleotides (ASOs) or antibody-based therapies. Extracellular protein aggregates might transmit to adjacent cells and thus artificially exported protein aggregates by an ATS would need to be handled further, like degradation by macrophages or excretion from the organism. Another potential therapeutic possibility could be an artificial pre-seeding of mHtt inclusions. In that way, an aggregation seed is generated on which mHtt oligomers could be sequestered. It has been proposed that small mHtt oligomers are the actual toxic species rather than the large inclusions (*67, 68*). These toxic, small oligomers could potentially be sequesterd in large inclusions using an artificial pre-seeding approach. Interestingly, *C. elegans* possesses a mechanism to clear its neurons from protein aggregates, including mutant human Huntingtin via membrane-surrounded vesicles called exophers (*69*).

In conclusion, our work demonstrates that protein aggregate inclusions are amenable to several kinds of spatial manipulations with potential therapeutic opportunity in humans. In addition, the here described artificial aggregate targeting system has the potential to be useful as an alternative, complementary approach to study the role of sPQC in aging and neurodegenerative disease.

## Materials and Methods

### Strains, plasmids, and growth conditions

Plasmids and strains are listed in tables S3 and S4, respectively. The strains used in this study are derivatives of BY4741 (*MAT****a*** *his3Δ1 leu2Δ0 met15Δ0 ura3Δ0*). Cells were cultured at 30 °C in rich YPD medium, in complete synthetic medium (CSM) or in synthetic dropout (SD) media (Formedium). For galactose induction, cells were grown overnight in SD-Ura with 2% (w/v) raffinose and 0.2% (w/v) glucose. Subsequently, cells were diluted to an OD600 of 0.1 in SD-Ura with 1% (w/v) galactose and 1% (w/v) raffinose and grown for 4.5 hours. Deletion strains were from the Yeast Knockout Collection (*70*). Strains with GFP-labeled endogenous proteins were from the yeast GFP clone collection (ThermoFisher Scientific) (*52*). All plasmids were verified by DNA sequencing. p416 25Q GPD was a gift from Susan Lindquist (Addgene plasmid # 1177 ; http://n2t.net/addgene:1177 ; RRID:Addgene_1177). p416 103Q GPD was a gift from Susan Lindquist (Addgene plasmid # 1180 ; http://n2t.net/addgene:1180 ; RRID:Addgene_1180). pAG416GPD-EGFP was a gift from Susan Lindquist (Addgene plasmid # 14196 ; http://n2t.net/addgene:14196 ; RRID:Addgene_14196). Human codon-optimized Hsp104 was generated through gblocks gene fragments (Integrated DNA Technologies).

### Strain construction

Transformations were done using the standard lithium acetate protocol and transformants were confirmed using PCR and phenotypic assays. Variants of ATS were genomically integrated at the MET15 locus, by cutting the plasmids containing the ATS variant with PmeI (ThermoFisher Scientific), which introduces two cuts and exposes homology regions to the *MET15* locus. The integrated ATS variants contained the *CYC1* terminator and were under the control of the weak constitutive *ADH1* promoter, unless stated otherwise.

### Growth time analysis

Yeast cells were grown overnight at 30 °C, diluted to an optical density at 600 nm (OD600) of 0.1 and distributed in a 24-well microtiter plate with a total volume of 800 µl per well. The microtiter plate was then sealed with a MicroAmp optical adhesive film (ThermoFisher Scientific) and punctured with a 23 Gauge needle for air exchange. A plate reader (Biotek Synergy 2 SL Microplate Reader Luminescence and Biotek Gen5 Data Analysis Software) was used to monitor OD600 of the cultures every 15 minutes for 24 hours. For experiments with inducible expression, the galactose induction was started at the beginning of the plate reader monitoring.

### Serial growth assay

Exponentially growing cells were cultured in YPD and diluted to an optical density of 0.1 followed by 10-fold dilutions. 5 µl of each dilution was spotted onto solid YPD agar media. Images were taken after 48 h of growth at the indicated temperature.

### Human cell culture and transient transfection

Human HEK293 cells were cultured at 37 °C and 5% CO_2_ in DMEM (ThermoFisher Scientific), supplemented with 10% fetal bovine serum (ThermoFisher Scientific), 100 units/mL penicillin and 100 µg/mL streptomycin (ThermoFisher Scientific). For transient transfection, 50,000 cells were seeded per well (24 well plate, 500 µl growth medium). After 24 h, cells were transfected with Lipofectamine 3000 (ThermoFisher Scientific) according to the manufacturer’s instruction using 3 µl transfection reagent per µg DNA and 2 µl of p3000 reagent per µg DNA. Per well 500 ng of DNA were used. After 24 h, cells were fixed in 3.7% formaldehyde solution and mounted using Fluoromount-G Mounting Medium, with DAPI (ThermoFisher Scientific).

### Fluorescence microscopy

Images were mostly obtained using an Axio Observer Z1 fluorescence microscope (Zeiss), equipped with an Axiocam 506 mono camera and a 20x, as well as a 100x oil objective lens (Plan-APOCHROMAT 100x/1.4 Oil DIC) controlled by the Zen blue software. Raw data were collected as Z-stacks and projected using ImageJ (NIH) with manual quantification.

### Time lapse microscopy

For yeast cells, SD medium containing agar was poured into a Gene Frame (ThermoFisher Scientific) attached to a cover glass. Cells were loaded and the agar pad was sealed with a coverslip. The cells were transferred into an incubation chamber maintained at 30 °C. Images were obtained using an Axio Observer 7 fluorescence microscope (Zeiss), equipped with an Axiocam 506 mono camera and a 20x, as well as 100x oil objective lens (Plan-APOCHROMAT 100x/1.4 Oil DIC). Focus stabilization was done with the Definite Focus module (Zeiss).

For human HEK293 cells, 40,000 cells were seeded into µ-Slide 4 Well chambered coverslips (Ibidi) using medium without phenol red. Cell transfection was performed as described above using medium without phenol red. After 24 h, HEPES pH 7.4 buffer was added to a final concentration of 25 mM and the cells were transferred to a incubation chamber maintained at 37 °C. Images were obtained using an Axio Observer 7 fluorescence microscope (Zeiss), equipped with an Axiocam 506 mono camera and a 20x objective lens.

### Heat shock treatments

Yeast cells were subjected to a continuous 38 °C heat shock in a shaking water bath, using 100 ml Erlenmeyer flasks.

### Hsf1 activity reporter assay

The activity of the Hsf1-dependent stress response was determined as described previously (*43*). Briefly, yeast cells were transformed with a plasmid (pAM10) carrying the luciferase NanoLuc under the control of a heat shock promoter (HSE) or with a control plasmid (pAM09). Nano-Glo substrate (Promega) was diluted 1:100 with the supplied lysis buffer and mixed 1:10 with cells grown in SD-Ura medium in a white 96-well plate. Bioluminescence was determined immediately, using a plate reader (Biotek Synergy 2 SL).

### Synthetic genetic array screen and high content microscopy

A genome-wide SGA screen was performed as described previously (*71*), using a BM3-BC colony handling robot (S&P Robotics Inc.). The screen was run in 1536 spot format with the GBP-Pea2 query strain (with expression of mHtt103QP-mCherry) in the Y7039 background, crossed to the yeast GFP clone collection (ThermoFisher Scientific) (*52*). Final strains were grown in 96 well plates in liquid SD medium to an OD600 of about 0.5. Cells were then transferred to a black 384 well plate with a glass bottom (Matriplate). DAPI was added to a final concentration of 3 µg/ml to visualize nuclei and whole cells. Imaging was then performed with a ImageXpress Micro high content microscope (Molecular Devices). The ratio of GFP intensity in the artificial inclusion in relation to the cytoplasm was measured, as well as the fraction of cells with artificial inclusion. From the initial screen the top 78 strains were selected (ratio × fraction, cutoff 0.55) and three more independent experiments were performed with those to validate the screen results. As controls, the corresponding strains from the GFP clone collection were used without GBP-Pea2 and mHtt103QP-mCherry. GO analysis of biological process enrichment was performed with the DAVID Functional Annotation Tool (david.ncifcrf.gov), using the GOTERM_BP_DIRECT option. For all GO analyses, default settings were used, and categories with a p-value of < 0.05 were considered as significantly enriched. Only enriched pathways with at least 3 screen hit genes were considered. Network analysis was performed by the STRING database (string-db.org) with all default setting but without text-mining. The STRING database creates a network from a gene list, based on its physical interactions and functional associations. The results were then exported to the cytoscape software (version 3.9.0) for visualization.

### Lifespan analysis by microdissection

Yeast replicative lifespan was measured by using Singer MSM micromanipulator and following standard procedures (*72*). Lifespan figures in main text depict a representative experiment. All lifespans were tested in at least two separate sets (n) of experiments. Lifespan p-values were determined using the Wilcoxon rank-sum test.

### Lifespan analysis by microfluidics

Microfluidics time-lapse microscopy were performed using a Zeiss Axio Observer 7 inverted fluorescence microscope with Definite Focus, equipped with an Axiocam 506 mono camera (Zeiss). Images were taken with a 40X objective. During the experiment, the microfluidics device (iBiochips) was heated to 30 °C in a heating chamber. The microscope was programmed to acquire brightfield images every 15 min for a total of about 65 hours. Occasionally, GFP images were also taken. Camera binning was set to 3. A stack of three planes was taken. Cells were grown overnight in YPD medium (2% glucose). On the next day, cells were diluted to an OD_600_ of 0.1 and grown until an OD of 0.6. Cells were put into a 1-ml syringe and four yeast strains were loaded at the same time into the microfluidics chip using a NE-4000 syringe pump (New Era Pumps) with a flow rate of 1 µl/min. As soon as the traps were filled, the cell loading ports were closed with a metal pin. The medium flow rate was set to 10 µl/min using a NE-1000 syringe pump (New Era Pumps). Sterile filtered YPD medium in a 60-ml syringe was used. Time-lapse images were analyzed manually. Amount of cell divisions were counted. Yeast RLS p-values were determined using the Gehan-Breslow-Wilcoxon test.

### Statistical analysis

Quantification and statistical analysis were performed using GraphPad Prism 9. Data in graphs are presented as mean +/- SEM and were analyzed using unpaired two-tailed t-test, if not stated otherwise. Growth curves were analyzed by two-way ANOVA. Replicative lifespan data was analyzed as described above. Significance levels * < 0.05, ** < 0.01, *** < 0.001, **** < 0.0001.

## Acknowledgments

We thank all members of the Nyström laboratory as well as Mikael Molin and Claes Andréasson for discussion; Harm Kampinga for the pHttQ119-EYFP plasmid; John McCullough for the pGag_eGFP plasmid; Lisa Larsson Berglund for yeast mHtt plasmids; Per Widlund for a MET15 locus integrative plasmid backbone (pPW411), for a Myo2 plasmid, as well as for pro3-1, guk1-7 and gus1-3 misfolding reporter plasmids and strains. Claes Andréasson for plasmids pAM09 and pAM10 for the Hsf1 activity reporter assay; Sarah Hanzén for mutant Tsa1 strains; Martí Aldea for the YDJ1-GFP-FS strain; Benedict Tan and Per Widlund for critically reading the manuscript.

## Funding

The study was supported by a research fellowship to A.F. from the German research foundation (DFG), as well as grants from the VR and Knut and Alice Wallenberg Foundation to T.N.

## Author contributions

Conceptualization, A.F., T.N.; Methodology, A.F., X.H.; Investigation, A.F., A.J., K.S.; Writing - Original Draft, A.F., T.N.; Writing - Review &Editing, A.F., T.N., A.J., K.S., X.H.; Supervision, Project Administration and Funding Acquisition, T.N., A.F.

## Competing interests

The authors declare that they have no competing interests.

## Data and materials availability

All data needed to evaluate the conclusions in the paper are present in the paper and/or the Supplementary Materials. Additional data related to this paper may be requested from the authors.

## Supplementary Materials

**Fig. S1.**
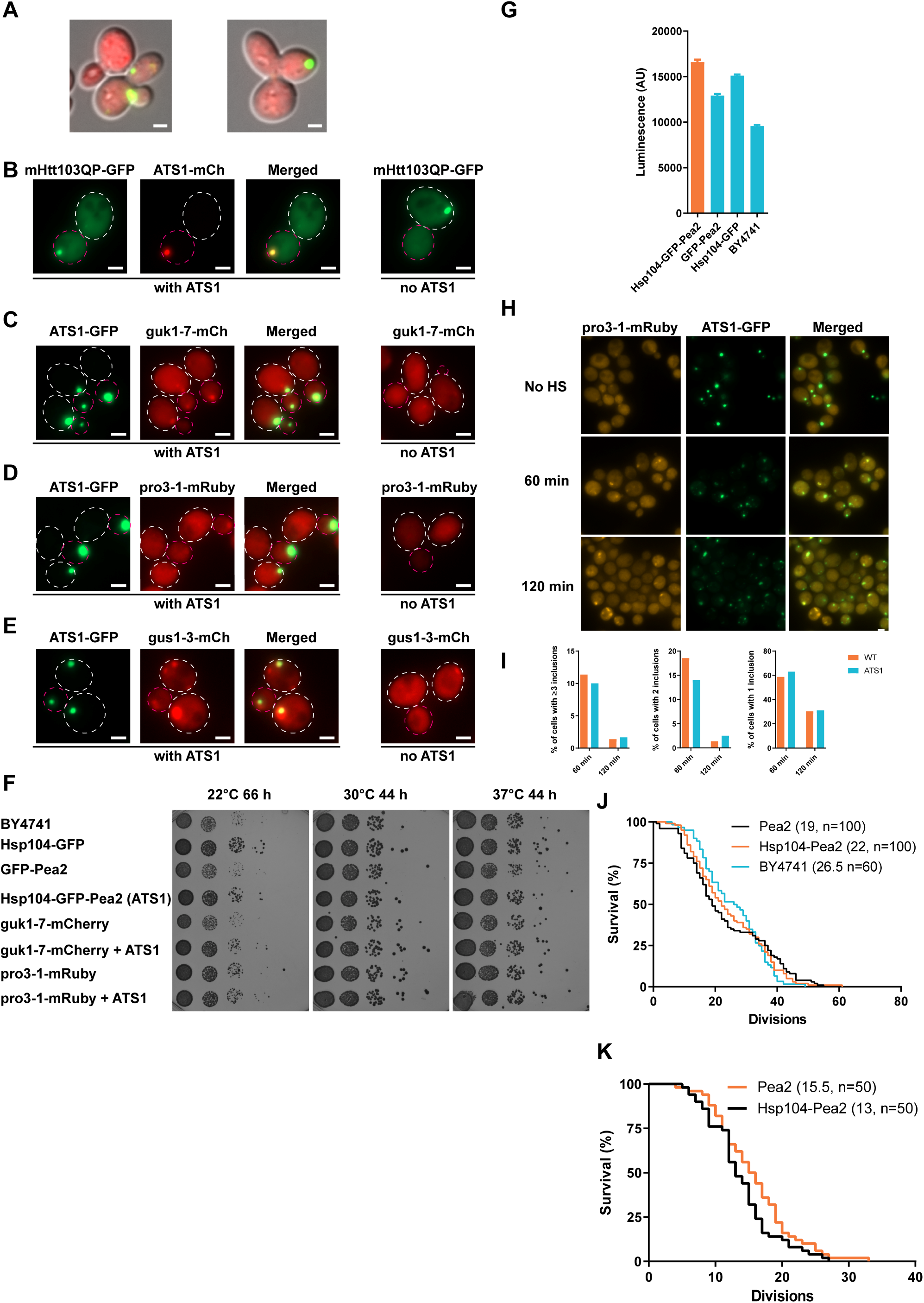
Generation of an aggregate targeting system to the bud. (**A**) Two representative fluorescence microscopy images showing cell division defects using Hsp104-GFP-Myo2 (green). This fusion protein was not used in later experiments. (**B**) Left panel: Representative fluorescence microscopy images of mHtt103QP-GFP aggregate transport into buds with ATS1. mHtt103QP-GFP was expressed constitutively (GPD promoter). Right panel: mHtt103QP aggregation in the absence of ATS1. Scale bar represents 2 µm. (**C**) Representative fluorescence microscopy images of guk1-7-mCherry (temperature-sensitive allele of *GUK1*) aggregate transport into buds with ATS1. The right panel is showing guk1-7-mCherry expression in the absence of ATS1. No heat shock was applied. (**D**) Representative fluorescence microscopy images of pro3-1-mRuby (temperature-sensitive allele of *PRO3*) aggregate transport into buds with ATS1. The right panel is showing pro3-1-mRuby expression in the absence of ATS1. No heat shock was applied. (**E**) Representative fluorescence microscopy images of gus1-3-mCherry (temperature-sensitive allele of *GUS1*) aggregate transport into buds with ATS1. The right panel is showing gus1-3-mCherry expression in the absence of ATS1. No heat shock was applied. (**F**) Serial growth assay of the indicated yeast strains at 22, 30 or 37 °C. (**G**) Bioluminescent determination of Hsf1 activity. n = 3 technical replicates. (**H**) Representative fluorescence microscopy images of pro3-1-mRuby disaggregation at 38 °C in the presence of ATS1-GFP. (**I**) Quantification of H. (**J**) Replicative lifespan analysis via microdissection of cells constitutively expressing ATS1 (*ADH1* promoter). Median lifespan is indicated in parentheses. N = 60-100 cells. (**K**) Replicative lifespan analysis of cells constitutively expressing ATS1, determined by microfluidics methodology. Median lifespan is indicated in parentheses. n = 50 cells. Scale bars represent 2 µm.

**Fig. S2.**
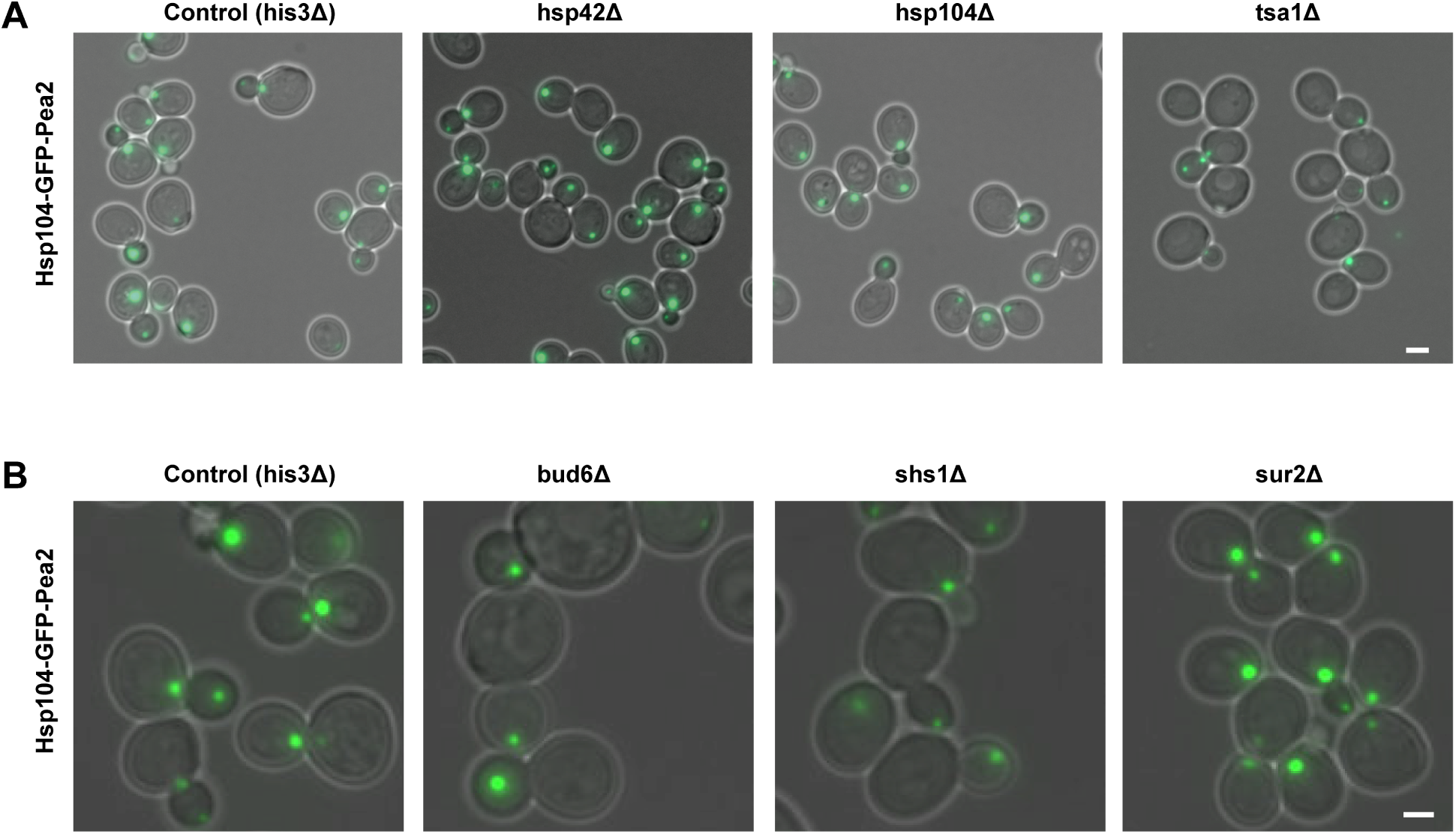
Genes and genetic modifications affecting ATS1. (**A**) ATS1 in deletion mutants of proteostasis genes. (**B**) ATS1 in deletion mutants of diffusion barrier genes. Scale bars represent 2 µm.

**Fig. S3.**
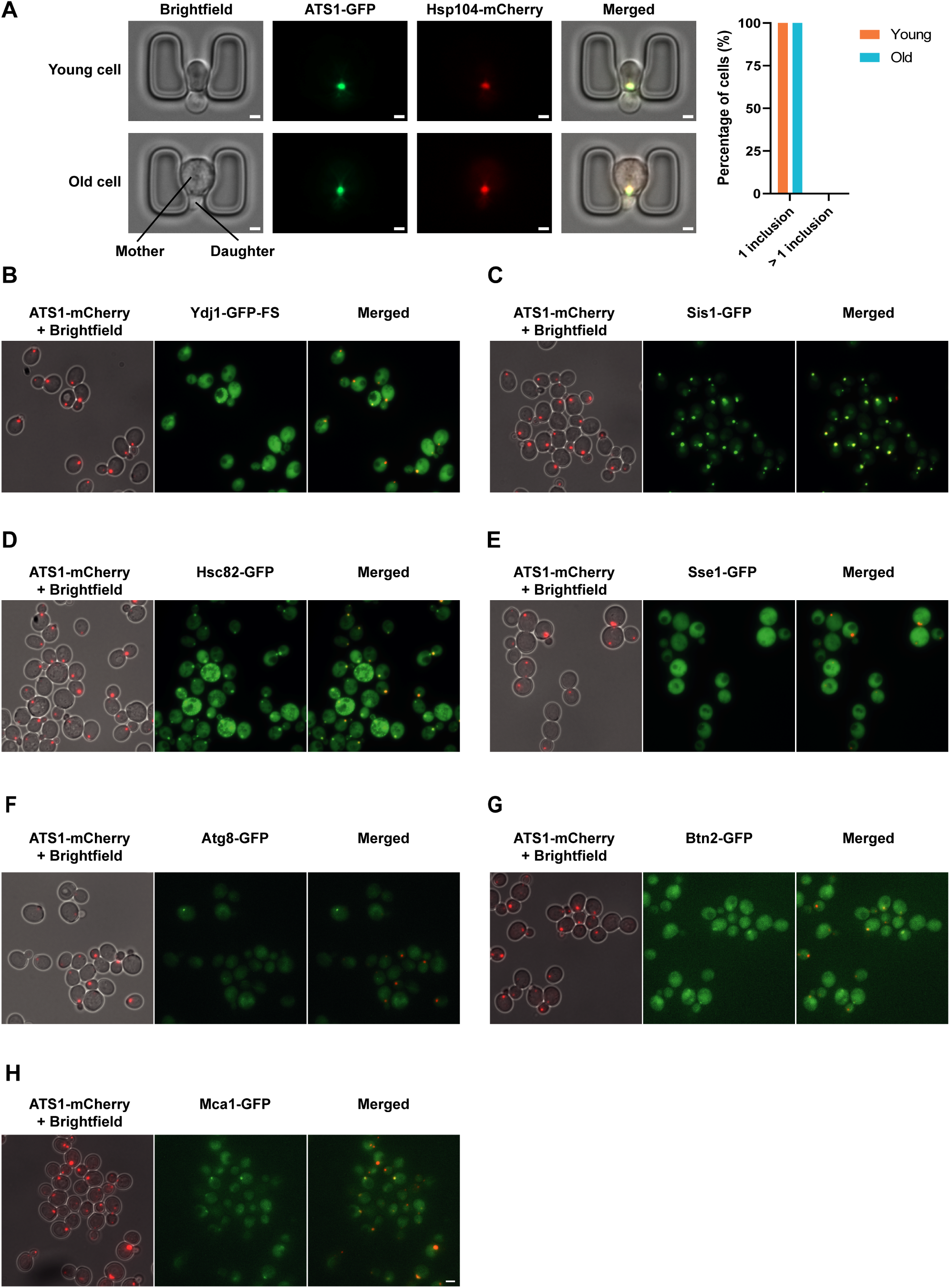
Endogenous proteins co-localizing with ATS1. (**A**) Representative fluorescence microscopy images of a young and an old cell (15 cell divisions) in a microfluidic chip, expressing Hsp104-GFP-Pea2 (ATS1-GFP). Endogenous Hsp104 was tagged with mCherry. The right panel shows a quantification of the amount of Hsp104-mCherry inclusions per cell (n = 50 cells). (**B**) Co-localization of ATS1 (red, mCherry-labeled) with endogenous Ydj1 (green, GFP label located between the J domain and the FS domain). (**C**) Co-localization of ATS1 (red, mCherry-labeled) with endogenous Sis1 (green, GFP-labeled). (**D**) Co-localization of ATS1 (red, mCherry-labeled) with endogenous Hsc82 (green, GFP-labeled). (**E**) No co-localization of ATS1 (red, mCherry-labeled) with endogenous Sse1 (green, GFP-labeled). (**F**) No co-localization of ATS1 (red, mCherry-labeled) with endogenous Atg8 (green, GFP-labeled). (**G**) Co-localization of ATS1 (red, mCherry-labeled) with endogenous Btn2 (green, GFP-labeled) only after a heat shock (110 min 38 °C). (**H**) Co-localization of ATS1 (red, mCherry-labeled) with endogenous Mca1 (green, GFP-labeled) only after a heat shock (110 min 38 °C). Scale bars represent 2 µm.

**Fig. S4.**
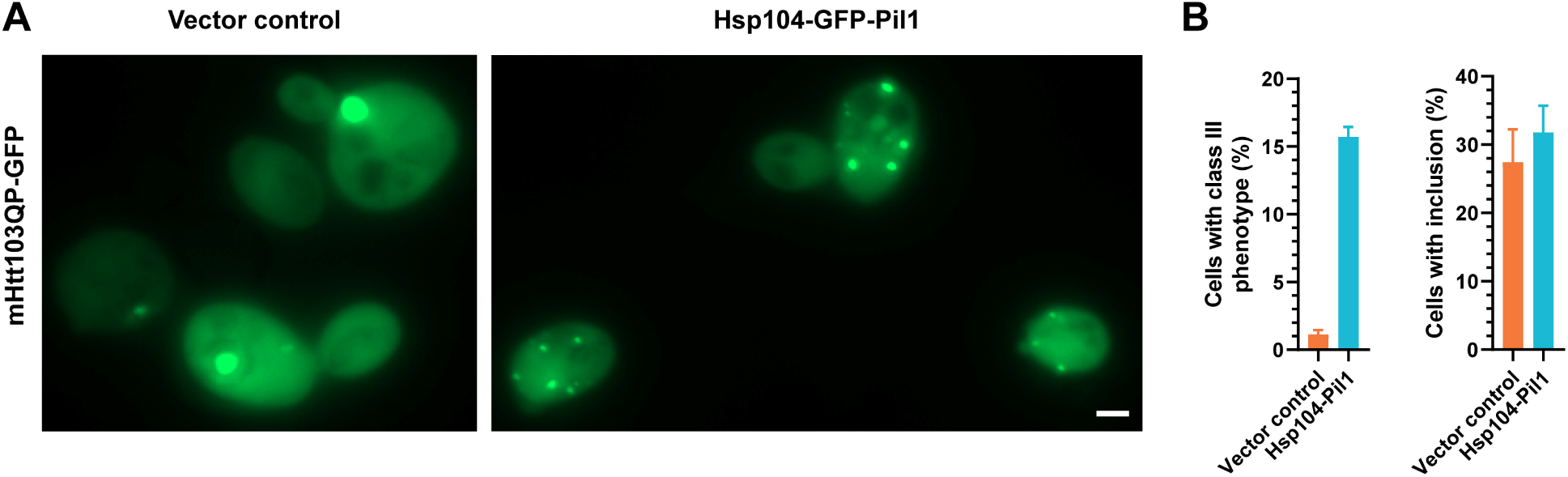
Manipulating aggregate inclusion site location to other sub-cellular locations. (**A**) Representative fluorescence microscopy images showing artificial targeting of Htt103QP-GFP aggregates to eisosomes by using Hsp104-GFP-Pil1. The expression of mHtt103QP-GFP was under the control of the constitutive GPD promoter. (**B**) Quantification of the amount of mHtt103QP-GFP aggregate inclusions per cell. n = 3 independent experiments. Class III phenotype: 3 or more aggregate inclusions per cell. Scale bar represents 2 µm.

**Fig. S5.**
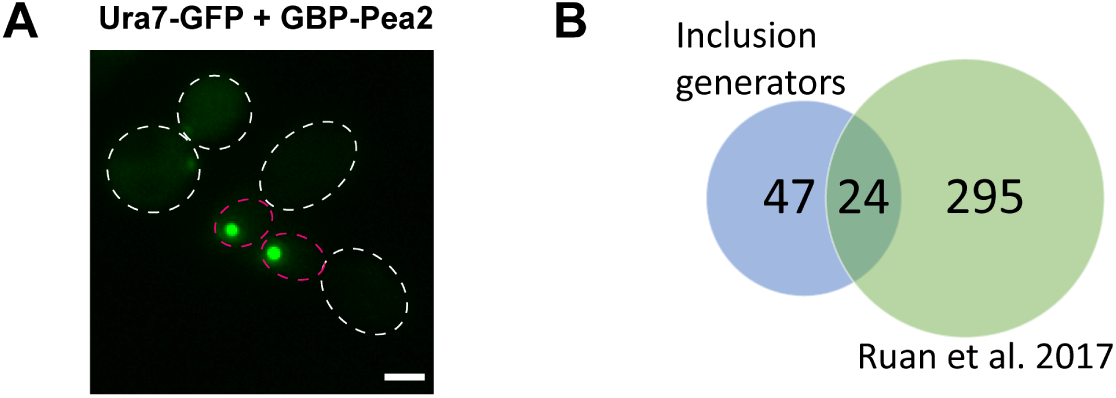
Generation of different types of artificial inclusions by generating Pea2 chimeras. (**A**) Representative fluorescence microscopy image of Ura7-GFP from inclusion generator screen (i.e. in the presence of GBP-Pea2). Scale bar represents 2 µm. (**B**) Venn diagram of the inclusion generator screen hits and proteins found in a screen for heat-shock-induced aggregates from Ruan et al. 2017.

**Table S1**.

Screen results for searching for other inclusion generators (IGs) than Hsp104.

**Table S2**.

Enrichment analysis of inclusion generators (IGs) for oligomerization.

**Table S3**.

Plasmid list.

**Table S4**.

Yeast strain list.

**Movie S1**.

Transport of mHtt103QP-GFP into the daughter cell using ATS1.

## Notes

### Competing Interest Statement

The authors have declared no competing interest.

