## Supplementary material for "Artificial Hsp100-mediated systems for re-localizing protein aggregates": Table S3 and S4

**Supplementary table 3. Plasmid list.**

| **Plasmid** | **Replicon** | **Promoter** | **Gene** | **Backbone** | **Parent** | **Selection marker** | **Reference** |
| --- | --- | --- | --- | --- | --- | --- | --- |
| p416 GPD | CEN/ARS | GPD | GFP | pRS416 | p416 GPD | URA3 | Krobitsch and Lindquist (2000) |
| p416 25Q GPD | CEN/ARS | GPD | mHtt25QP-GFP | pRS416 | p416 GPD | URA3 | Krobitsch and Lindquist (2000) |
| p416 103Q GPD | CEN/ARS | GPD | mHtt103QP-GFP | pRS416 | p416 GPD | URA3 | Krobitsch and Lindquist (2000) |
| pAM09 | CEN/ARS | - | - |  |  | URA3 | Masser et al. (2016) |
| pAM10 | CEN/ARS | CYC1–HSE | NanoLuc |  |  | URA3 | Masser *et al.* (2016) |
| pAF039 | - | - | MET15 homology regions | pRS303 | pPW411 | G418 | This study |
| pAF044 | - | Tet3 | Hsp104-GFP-Myo2(AA1-1086) | pRS303 | pAF039 | G418 | This study |
| pAF063 | - | ADH1 | Hsp104-GFP-Pea2 | pRS303 | pAF039 | G418 | This study |
| pAF066 | - | ADH1 | GFP-Pea2 | pRS303 | pAF039 | G418 | This study |
| pAF067 | - | ADH1 | Hsp104-GFP | pRS303 | pAF039 | G418 | This study |
| pAF070 | - | ADH1 | Hsp104-GFP-Pea2 | pRS303 | pPW411 | NAT | This study |
| pAF071 | - | ADH1 | Hsp104-GFP-Snf7 | pRS303 | pAF039 | G418 | This study |
| pAF075 | - | ADH1 | Hsp104-mCherry-Pea2 | pRS303 | pAF039 | G418 | This study |
| pAF083 | - | ADH1 | Hsp104(Δ22)-GFP-Pea2 | pRS303 | pAF063 | G418 | This study |
| pAF084 | - | ADH1 | Hsp104(Δ38)-GFP-Pea2 | pRS303 | pAF063 | G418 | This study |
| pAF095 | - | GPD | Hsp104-GFP-Pil1 | pRS303 | pAF039 | G418 | This study |
| pAF099 | - | ADH1 | Hsp104(Δ38, ΔATP1)-GFP-Pea2 | pRS303 | pAF084 | G418 | This study |
| pAF100 | - | ADH1 | Hsp104(Δ38, ΔATP2)-GFP-Pea2 | pRS303 | pAF084 | G418 | This study |
| pAF103 | - | ADH1 | Hsp104(ΔATP1)-GFP-Pea2 | pRS303 | pAF063 | G418 | This study |
| pAF104 | - | ADH1 | Hsp104(ΔATP2)-GFP-Pea2 | pRS303 | pAF063 | G418 | This study |
| pAF107 | - | ADH1 | Hsp104-GFP-p53tet | pRS303 | pAF039 | G418 | This study |
| pAF111 | - | GPD | Hsp104-Pil1 | pRS303 | pAF095 | G418 | This study |
| pAF112 | - | ADH1 | Hsp104(ΔATP1, ΔATP2)-GFP-Pea2 | pRS303 | pAF104 | G418 | This study |
| pAF118 | - | ADH1 | GBP-Pea2 | pRS303 | pAF039 | G418 | This study |
| pAF130 | - | ADH1 | Tsa1(S78D)-GFP-Pea2 | pRS303 | pAF039 | G418 | This study |
| pAF131 | - | ADH1 | Hsp42(ΔCTD)-GFP-Pea2 | pRS303 | pAF039 | G418 | This study |
| pAF133 | - | Tet1 | Hsp104-GFP-Pea2 | pRS303 | pAF039 | G418 | This study |
| pAF136 | - | ADH1 | Tsa1(ΔOI1)-GFP-Pea2 | pRS303 | pAF039 | G418 | This study |
| pAF137 | - | ADH1 | Tsa1(ΔOI2)-GFP-Pea2 | pRS303 | pAF039 | G418 | This study |
| pAF139 | - | ADH1 | Hsp42(ΔACD)-GFP-Pea2 | pRS303 | pAF039 | G418 | This study |
| pAF142 | - | Tet1 | GFP-Pea2 | pRS303 | pAF039 | G418 | This study |
| pAF145 | - | ADH1 | Hsp42(ΔACD, ΔCTD)-GFP-Pea2 | pRS303 | pAF039 | G418 | This study |
| pAF149 | - | ADH1 | Hsp42(ΔNTD)-GFP-Pea2 | pRS303 | pAF039 | G418 | This study |
| pAF154 | - | ADH1 | Hsp42(CTD)-GFP-Pea2 | pRS303 | pAF039 | G418 | This study |
| pAF155 | - | ADH1 | Hsp42(ACD)-GFP-Pea2 | pRS303 | pAF039 | G418 | This study |
| pAF171 | - | Tet1 | Hsp104-GFP | pRS303 | pAF039 | G418 | This study |
| pAF168 | SV40 | CMV | mCherry-Gag | pcdna4 |  | NeoR/Amp | This study |
| pAF180 | SV40 | CMV | Gag | pcdna4 |  | NeoR/Amp | This study |
| pAF181 | SV40 | CMV | Hsp104(codon optimized)-mCherry-Gag | pcdna4 |  | NeoR/Amp | This study |
| pAF195 | SV40 | CMV | Hsp104(codon optimized)-Gag | pcdna4 | pAF181 | NeoR/Amp | This study |
| pGag_eGFP | SV40 | CMV | Gag from HIV | pEGFP-N1 |  | NeoR/KanR | Hermida-Matsumoto and Resh (2000) |
| pHttQ119-EYFP | SV40 | CMV | mHtt119Q | pEYFP-N1 |  | NeoR/KanR | Rujano et al. (2006) |
| pPW351 | - | GPD | pro3-1-GFP | pRS403 |  | HIS3 | Schneider et al. (2018) |
| pPW390 | - | GPD | pro3-1-mRuby2 | pRS405 |  | LEU2 | Schneider *et al.* (2018) |
| pPW411 | - | - | MET15 homology regions | pRS303 |  | NAT | This study |
| pRS413-MYO2 | CEN/ARS | MYO2 | MYO2 | pRS413 |  | HIS3 | Catlett and Weisman (1998) |
| pRS416 | CEN/ARS | - | - | pRS416 |  | URA3 | Sikorski and Hieter (1989) |
| pRS416::GPDp-Htt103QP-mCherry | CEN/ARS | GPD | mHtt103QP-mCherry | pRS416 | p416 GPD | URA3 | This study |
| pYES2- GFP | 2µ | GAL1 | GFP | pYES2 |  | URA3 | Prévéral et al. (2006) |
| pYES2-Htt25Q-GFP | 2µ | GAL1 | mHtt25Q-GFP | pYES2 |  | URA3 | Meriin et al. (2007) |
| pYES2-Htt25QP-GFP | 2µ | GAL1 | mHtt25QP-GFP | pYES2 |  | URA3 | Meriin *et al.* (2007) |
| pYES2-Htt103Q-GFP | 2µ | GAL1 | mHtt103Q-GFP | pYES2 |  | URA3 | Meriin *et al.* (2007) |
| pYES2-Htt103QP-GFP | 2µ | GAL1 | mHtt103QP-GFP | pYES2 |  | URA3 | Meriin *et al.* (2007) |
| pYES2-Htt25Q- mCherry | 2µ | GAL1 | mHtt25Q-mCherry | pYES2 |  | URA3 | Meriin *et al.* (2007) |
| pYES2-Htt25QP- mCherry | 2µ | GAL1 | mHtt25QP-mCherry | pYES2 |  | URA3 | Meriin *et al.* (2007) |
| pYES2-Htt103Q- mCherry | 2µ | GAL1 | mHtt103Q-mCherry | pYES2 |  | URA3 | Meriin *et al.* (2007) |
| pYES2-Htt103QP- mCherry | 2µ | GAL1 | mHtt103QP-mCherry | pYES2 |  | URA3 | Meriin *et al.* (2007) |

**Supplementary table 4. Yeast strain list.**

| **ID** | **Name** | **Genotype** | **Parent strain** | **Ingegrated plasmid** | **Reference** |
| --- | --- | --- | --- | --- | --- |
| BY4741 | BY4741 | *MATa his3∆1 leu2∆0 met15∆0 ura3∆0* |  |  |  |
| guk1-7-mCherry | guk1-7-mCherry | *MATa his3∆1 leu2∆0 ura3∆0 met15∆0 lys2::GPDp-guk1-7-mCherry-PGK1t-LEU2* |  |  | Schneider *et al.* (2018) |
| PW1339 | gus1-3-mCherry | *MATa his3∆1 leu2∆0 ura3∆0 met15∆0 lys2::GPDp-gus1-3-mCherry-PGK1t-LEU2* |  |  | Schneider *et al.* (2018) |
| R1158 | CMV-tTA | URA3::CMV-tTA MATa his3Δ1 leu2Δ0 met15∆0 |  |  | Mnaimneh et al. (2004) |
| SMH223 | ssa1∆ ssa2∆ | MATa ssa1Δ::hphMX ssa2Δ::kanMX4 his3Δ1 LYS2+ leu2Δ0 met15Δ0 ura3Δ0 |  |  | Hill et al. (2014) |
| SH164 | Tsa1-GFP | MAT a ura3-52 leu2Δ1 trp1Δ63 his3Δ200 lys2ΔBgl hom3-10, ade2Δ1, ade8, hxt13Δ::URA3 TSA1(WT)_GFP::NAT/TRP1 met15Δ0 |  |  | Hanzen et al. (2016) |
| SH165 | Tsa1_C48S-GFP | MAT a ura3-52 leu2Δ1 trp1Δ63 his3Δ200 lys2ΔBgl hom3-10, ade2Δ1, ade8, hxt13Δ::URA3 tsa1C48S-GFP::NAT/TRP1 cyh2 met15Δ0 |  |  | Hanzen *et al.* (2016) |
| SH166 | Tsa1_C171S-GFP | MAT a ura3-52 leu2Δ1 trp1Δ63 his3Δ200 lys2ΔBgl hom3-10, ade2Δ1, ade8, hxt13Δ::URA3 tsaC171S-GFP::NAT/TRP1 cyh2 met15Δ0 |  |  | Hanzen *et al.* (2016) |
| SH169 | Tsa1_C48,171S-GFP | MAT a ura3-52 leu2Δ1 trp1Δ63 his3Δ200 lys2ΔBgl hom3-10, ade2Δ1, ade8, hxt13Δ::URA3 tsa1C48S,C171S-GFP::NAT/TRP1 cyh2 met15Δ0 |  |  | Hanzen *et al.* (2016) |
| SH192 | Tsa1-GFP | MATα, his3Δ1, leu20, lysΔ0, ura3Δ0 Tsa1-GFP::hph met15Δ0 |  |  | Hanzen *et al.* (2016) |
| SH193 | Tsa1_DYF-GFP | MATα, his3Δ1, leu20, lysΔ0, ura3Δ0 Tsa1DYF-GFP::hph met15Δ0 |  |  | Hanzen *et al.* (2016) |
| yAF023 | pro3-1-GFP | *MAT***a** *Hsp104Δ::kanMX4* *his3Δ1::HIS3-pGPD-pro3-1-GFP-tPgk1 leu2Δ0 met15Δ0 ura3Δ0* | BY4741 | pPW351 | This study |
| yAF100 | Hsp104-GFP-Myo2(1-1086) | *MATa his3∆1 leu2∆0 met15∆0 ura3∆0 Gal4::His3-pMyo2-GEV-tPgk1 Ade4::NatMX-rtAct1-pGal1-(rtTA-SE-G72P)-tPgk1 LYS2::Leu2-pGPD-pro3-1-mRuby Met15::TET3p-Hsp104-GFP-Myo2(1-1086)-CYC1t-KanMX* | yAF049 | pAF044 | This study |
| yAF204 | ADH1p-Hsp104-GFP-Pea2 | *MATa his3∆1 leu2∆0 ura3∆0 met15∆0::ADH1p-Hsp104-GFP-Pea2-CYC1t-KanMX* | BY4741 | pAF063 | This study |
| yAF217 | Hsp104-mCherry + Hsp104-GFP-Pea2 | *MATa his3∆1 leu2∆0 ura3∆0 HSP104-mCherry-hphNT1 met15∆0::ADH1p-Hsp104-GFP-Pea2-CYC1t-KanMX* | BY4741 | pAF063 | This study |
| yAF219 | ADH1p-Hsp104-GFP-Pea2 + Hsp42-mRuby | *MATa his3∆1 leu2∆0 ura3∆0 met15∆0::ADH1p-Hsp104-GFP-Pea2-CYC1t-KanMX HSP42-mRuby-hphNT1* | yAF204 |  | This study |
| yAF221 | Hsp104-GFP-Pea2 + pro3-1-mRuby2 | *MATa his3∆1 leu2∆0 ura3∆0 met15∆0::ADH1p-Hsp104-GFP-Pea2-CYC1t-KanMX lys2::GPDp-pro3-1-mRuby2-PGK1t-LEU2* | yAF204 |  | This study |
| yAF227 | ADH1p-Hsp104-GFP-Pea2 + pYES-GAL1p-Htt103QP-mCherry | *MATa his3∆1 leu2∆0 ura3∆0 met15∆0::ADH1p-Hsp104-GFP-Pea2-CYC1t-KanMX pYES-GAL1p-Htt103QP-mCherry(URA)* | yAF204 |  | This study |
| yAF229 | guk1-7-mCherry + ADH1p-Hsp104-GFP-Pea2 | *MATa his3∆1 leu2∆0 ura3∆0 met15∆0::ADH1p-Hsp104-GFP-Pea2-CYC1t-KanMX lys2::GPDp-guk1-7-mCherry-PGK1t-LEU2* | guk1-7-mCherry |  | This study |
| yAF235 | GPDp-pro3-1-mRuby2 | *MATa his3∆1 leu2∆0 ura3∆0 met15∆0 lys2::GPDp-pro3-1-mRuby2-PGK1t-LEU2* | BY4741 | pPW390 | This study |
| yAF241 | ADH1p-GFP-Pea2 | *MATa his3∆1 leu2∆0 ura3∆0 met15∆0::ADH1p-GFP-Pea2-CYC1t-KanMX* | BY4741 | pAF066 | This study |
| yAF242 | ADH1p-Hsp104-GFP | *MATa his3∆1 leu2∆0 ura3∆0 met15∆0::ADH1p-Hsp104-GFP-CYC1t-KanMX* | BY4741 | pAF067 | This study |
| yAF250 | ADH1p-Hsp104-GFP-Pea2 + pAM09 | *MATa his3∆1 leu2∆0 ura3∆0 met15∆0::ADH1p-Hsp104-GFP-Pea2-CYC1t-KanMX pAM09(URA)* | yAF204 |  | This study |
| yAF251 | ADH1p-Hsp104-GFP-Pea2 + pAM10 | *MATa his3∆1 leu2∆0 ura3∆0 met15∆0::ADH1p-Hsp104-GFP-Pea2-CYC1t-KanMX pAM10(URA)* | yAF204 |  | This study |
| yAF252 | ADH1p-GFP-Pea2 + pAM09 | *MATa his3∆1 leu2∆0 ura3∆0 met15∆0::ADH1p-GFP-Pea2-CYC1t-KanMX pAM09(URA)* | yAF241 |  | This study |
| yAF253 | ADH1p-GFP-Pea2 + pAM10 | *MATa his3∆1 leu2∆0 ura3∆0 met15∆0::ADH1p-GFP-Pea2-CYC1t-KanMX pAM10(URA)* | yAF241 |  | This study |
| yAF254 | ADH1p-Hsp104-GFP + pAM09 | *MATa his3∆1 leu2∆0 ura3∆0 met15∆0::ADH1p-Hsp104-GFP-CYC1t-KanMX pAM09(URA)* | yAF242 |  | This study |
| yAF255 | ADH1p-Hsp104-GFP + pAM10 | *MATa his3∆1 leu2∆0 ura3∆0 met15∆0::ADH1p-Hsp104-GFP-CYC1t-KanMX pAM10(URA)* | yAF242 |  | This study |
| yAF256 | BY4741 + pAM09 | *MATa his3∆1 leu2∆0 ura3∆0 met15Δ0 pAM09(URA)* | BY4741 |  | This study |
| yAF257 | BY4741 + pAM10 | *MATa his3∆1 leu2∆0 ura3∆0 met15Δ0 pAM10(URA)* | BY4741 |  | This study |
| yAF260 | ADH1p-Hsp104-GFP-Pea2 | *MATa his3∆1 leu2∆0 ura3∆0 met15∆0::ADH1p-Hsp104-GFP-Pea2-CYC1t-NATMX* | BY4741 |  | This study |
| yAF261 | bud6Δ + ADH1p-Hsp104-GFP-Pea2 | *MAT***a** *bud6Δ::kanMX4* *his3Δ1 leu2Δ0 ura3Δ0 met15∆0::ADH1p-Hsp104-GFP-Pea2-CYC1t-NATMX* | bud6Δ | pAF070 | This study |
| yAF262 | shs1Δ + ADH1p-Hsp104-GFP-Pea2 | *MAT***a** *shs16Δ::kanMX4* *his3Δ1 leu2Δ0 ura3Δ0 met15∆0::ADH1p-Hsp104-GFP-Pea2-CYC1t-NATMX* | shs1Δ | pAF070 | This study |
| yAF263 | sur2Δ + ADH1p-Hsp104-GFP-Pea2 | *MAT***a** *sur2Δ::kanMX4* *his3Δ1 leu2Δ0 ura3Δ0 met15∆0::ADH1p-Hsp104-GFP-Pea2-CYC1t-NATMX* | sur2Δ | pAF070 | This study |
| yAF264 | hsp42Δ + ADH1p-Hsp104-GFP-Pea2 | *MAT***a** *hsp42Δ::kanMX4* *his3Δ1 leu2Δ0 ura3Δ0 met15∆0::ADH1p-Hsp104-GFP-Pea2-CYC1t-NATMX* | hsp42Δ | pAF070 | This study |
| yAF270 | ADH1p-Hsp104-GFP-Snf7 | *MATa his3∆1 leu2∆0 ura3∆0 met15∆0::ADH1p-Hsp104-GFP-Snf7-CYC1t-KanMX* | BY4741 | pAF071 | This study |
| yAF277 | gus1-3-mCherry + ADH1p-Hsp104-GFP-Pea2 | *MATa his3∆1 leu2∆0 ura3∆0 met15∆0::ADH1p-Hsp104-GFP-Pea2-CYC1t-KanMX lys2::GPDp-gus1-3-mCherry-PGK1t-LEU2* | PW1339 | pAF063 | This study |
| yAF280 | ADH1p-Hsp104-GFP-Snf7 + pYES-GAL1p-Htt103QP-mCherry | *MATa his3∆1 leu2∆0 ura3∆0 met15∆0::ADH1p-Hsp104-GFP-Snf7-CYC1t-KanMX pYES-GAL1p-Htt103QP-mCherry(URA)* | yAF270 |  | This study |
| yAF289 | Ssa1-GFP + ADH1p-Hsp104-mCherry-Pea2 | *MATa his3∆1 leu2∆0 ura3∆0 SSA1-GFP-HIS3 met15∆0::ADH1p-Hsp104-mCherry-Pea2-CYC1t-KanMX* | Ssa1-GFP | pAF075 | This study |
| yAF294 | pADH1-Hsp104(∆22)-GFP-Pea2 | *MATa his3∆1 leu2∆0 ura3∆0 met15∆0::ADH1p-HSP104(∆22)-GFP-PEA2-CYC1t-KanMX* | BY4741 | pAF083 | This study |
| yAF295 | pADH1-Hsp104(∆38)-GFP-Pea2 | *MATa his3∆1 leu2∆0 ura3∆0 met15∆0::ADH1p-HSP104(∆38)-GFP-PEA2-CYC1t-KanMX* | BY4741 | pAF084 | This study |
| yAF306 | GPDp-GFP + ADH1p-Hsp104-mCherry-Pea2 | MATa his3∆1::GPDp-GFP-HIS3 leu2∆0 ura3∆0 met15∆0::ADH1p-Hsp104-mCherry-Pea2-CYC1t-KanMX | GPDp-GFP | pAF075 | This study |
| yAF332 | GPDp-Hsp104-GFP-PIL1 | *MATa his3∆1 leu2∆0 ura3∆0 met15∆0::GPDp-HSP104-GFP-PIL1-CYC1t-KanMX* | BY4741 | pAF095 | This study |
| yAF334 | ADH1p-Hsp104(∆38∆ATP1)-eGFP-Pea2 | *MATa his3∆1 leu2∆0 ura3∆0 met15∆0::ADH1p-Hsp104(∆38∆ATP1)-eGFP-PEA2-CYC1t-KanMX* | BY4741 | pAF099 | This study |
| yAF335 | ADH1p-Hsp104(∆38∆ATP2)-eGFP-Pea2 | *MATa his3∆1 leu2∆0 ura3∆0 met15∆0::ADH1p-Hsp104(∆38∆ATP2)-eGFP-PEA2-CYC1t-KanMX* | BY4741 | pAF100 | This study |
| yAF346 | GPDp-Hsp104-GFP-PIL1 + Htt103QP-mCh. | *MATa his3∆1 leu2∆0 ura3∆0 met15∆0::GPDp-HSP104-GFP-PIL1-CYC1t-KanMX pYES2-GAL1p-Htt103QP-mCherry(URA)* | yAF332 |  | This study |
| yAF350 | Ydj1-GFP-FS + ADH1p-Hsp104-mCherry-Pea2 | MATa his3∆1 leu2∆0 met15∆0::ADH1p-Hsp104-mCherry-Pea2-CYC1t-KanMX ura3∆0 YDJ1-GFP-FS-HIS3 | Ydj1-GFP-FS | pAF075 | This study |
| yAF351 | Sis1-GFP + ADH1p-Hsp104-mCherry-Pea2 | MATa his3∆1 leu2∆0 met15∆0::ADH1p-Hsp104-mCherry-Pea2-CYC1t-KanMX ura3∆0 SIS1-GFP-HIS3 | Sis1-GFP | pAF075 | This study |
| yAF352 | Btn2-GFP + ADH1p-Hsp104-mCherry-Pea2 | MATa his3∆1 leu2∆0 met15∆0::ADH1p-Hsp104-mCherry-Pea2-CYC1t-KanMX ura3∆0 BTN2-GFP-HIS3 | Btn2-GFP | pAF075 | This study |
| yAF353 | Mca1-GFP + ADH1p-Hsp104-mCherry-Pea2 | MATa his3∆1 leu2∆0 met15∆0::ADH1p-Hsp104-mCherry-Pea2-CYC1t-KanMX ura3∆0 MCA1-GFP-HIS3 | Mca1-GFP | pAF075 | This study |
| yAF355 | ssa1∆ ssa2∆ + ADH1p-Hsp104-GFP-Pea2 | MATa ssa1Δ::hphMX ssa2Δ::kanMX4 his3Δ1 LYS2+ leu2Δ0 met15Δ0::ADH1p-Hsp104-GFP-Pea2-CYC1t-NATMX ura3Δ0 | SMH223 | pAF070 | This study |
| yAF357 | ADH1p-Hsp104-mCherry-Pea2 + GFP-Atg8 | *MATa his3∆1 leu2∆0 ura3∆0 met15∆0::ADH1p-Hsp104-mCherry-Pea2-CYC1t-KanMX GFP-ATG8-HIS3* | yAF274 | pAF075 | This study |
| yAF358 | ADH1p-Hsp104(∆ATP1)-eGFP-Pea2 | *MATa his3∆1 leu2∆0 ura3∆0 met15∆0::ADH1p-Hsp104(∆ATP1)-eGFP-PEA2-CYC1t-KanMX* | BY4741 | pAF103 | This study |
| yAF359 | ADH1p-Hsp104(∆ATP2)-eGFP-Pea2 | *MATa his3∆1 leu2∆0 ura3∆0 met15∆0::ADH1p-Hsp104(∆ATP2)-eGFP-PEA2-CYC1t-KanMX* | BY4741 | pAF104 | This study |
| yAF367 | ssa1∆ + ADH1p-Hsp104-GFP-Pea2 | *MAT***a** *ssa1Δ::kanMX4* *his3Δ1 leu2Δ0 met15Δ0::ADH1p-Hsp104-GFP-Pea2-CYC1t-NATMX ura3Δ0* | ssa1∆ | pAF070 | This study |
| yAF368 | ssa2∆ + ADH1p-Hsp104-GFP-Pea2 | *MAT***a** *ssa2Δ::kanMX4* *his3Δ1 leu2Δ0 met15Δ0::ADH1p-Hsp104-GFP-Pea2-CYC1t-NATMX ura3Δ0* | ssa2∆ | pAF070 | This study |
| yAF369 | hsp104∆ + ADH1p-Hsp104-GFP-Pea2 | *MAT***a** *hsp104Δ::kanMX4* *his3Δ1 leu2Δ0 met15Δ0::ADH1p-Hsp104-GFP-Pea2-CYC1t-NATMX ura3Δ0* | hsp104∆ | pAF070 | This study |
| yAF370 | npr3Δ + ADH1p-Hsp104-GFP-Pea2 | *MAT***a** *npr3Δ::kanMX4* *his3Δ1 leu2Δ0 met15Δ0::ADH1p-Hsp104-GFP-Pea2-CYC1t-NATMX ura3Δ0* | npr3Δ | pAF070 | This study |
| yAF371 | tco89Δ + ADH1p-Hsp104-GFP-Pea2 | *MAT***a** *tco89Δ::kanMX4* *his3Δ1 leu2Δ0 met15Δ0::ADH1p-Hsp104-GFP-Pea2-CYC1t-NATMX ura3Δ0* | tco89Δ | pAF070 | This study |
| yAF372 | his3Δ + ADH1p-Hsp104-GFP-Pea2 | *MAT***a** *his3Δ::kanMX4* *his3Δ1 leu2Δ0 met15Δ0::ADH1p-Hsp104-GFP-Pea2-CYC1t-NATMX ura3Δ0* | his3Δ | pAF070 | This study |
| yAF373 | Hsc82-GFP + ADH1p-Hsp104-mCherry-Pea2 | MATa his3∆1 leu2∆0 met15∆0::ADH1p-Hsp104-mCherry-Pea2-CYC1t-KanMX ura3∆0 HSC82-GFP-HIS3 | Hsc82-GFP | pAF075 | This study |
| yAF374 | Tsa1-GFP + ADH1p-Hsp104-mCherry-Pea2 | MATa his3∆1 leu2∆0 met15∆0::ADH1p-Hsp104-mCherry-Pea2-CYC1t-KanMX ura3∆0 SIS1-GFP-HIS3 | Tsa1-GFP | pAF075 | This study |
| yAF375 | Sse1-GFP + ADH1p-Hsp104-mCherry-Pea2 | MATa his3∆1 leu2∆0 met15∆0::ADH1p-Hsp104-mCherry-Pea2-CYC1t-KanMX ura3∆0 Sse1-GFP-HIS3 | Sse1-GFP | pAF075 | This study |
| yAF378 | ADH1p-Hsp104-GFP-p53TETD | *MATa his3∆1 leu2∆0 ura3∆0 met15∆0::ADH1p-Hsp104-GFP-p53TETD-CYC1t-KanMX* | BY4741 | pAF107 | This study |
| yAF385 | ADH1p-Hsp104-mCherry-Pea2 + pRS416::GPDp-Htt103QP-GFP | *MATa his3∆1 leu2∆0 ura3∆0 met15∆0::ADH1p-Hsp104-mCherry-Pea2-CYC1t-KanMX pRS416::GPDp-Htt103QP-GFP(URA)* | yAF274 |  | This study |
| yAF386 | met15∆ vector control + pRS416 | *MATa his3∆1 leu2∆0 ura3∆0 met15∆0::KanMX pRS416(URA)* | yAF290 |  | This study |
| yAF387 | met15∆ vector control + pRS416::GPDp-GFP | *MATa his3∆1 leu2∆0 ura3∆0 met15∆0::KanMX pRS416::GPDp-GFP(URA)* | yAF290 |  | This study |
| yAF388 | met15∆ vector control + pRS416::GPDp-Htt25Q-GFP | *MATa his3∆1 leu2∆0 ura3∆0 met15∆0::KanMX pRS416::GPDp-Htt25Q-GFP(URA)* | yAF290 |  | This study |
| yAF389 | met15∆ vector control + pRS416::GPDp-Htt103QP-GFP | *MATa his3∆1 leu2∆0 ura3∆0 met15∆0::KanMX pRS416::GPDp-GFP(URA)* | yAF290 |  | This study |
| yAF393 | GPDp-Hsp104-Pil1 (No GFP) | *MATa his3∆1 leu2∆0 ura3∆0 met15∆0::GPDp-HSP104-PIL1-CYC1t-KanMX* | BY4741 | pAF111 | This study |
| yAF394 | ADH1p-Hsp104∆ATP1,∆ATP2-eGFP-Pea2 | *MATa his3∆1 leu2∆0 ura3∆0 met15∆0::ADH1p-Hsp104∆ATP1,∆ATP2-eGFP-PEA2-CYC1t-KanMX* | BY4741 | pAF112 | This study |
| yAF411 | GPDp-Hsp104-Pil1 + pRS416::GPDp | *MATa his3∆1 leu2∆0 ura3∆0 met15∆0::GPDp-HSP104-PIL1-CYC1t-KanMX pRS416::GPDp-GFP(URA)* | yAF393 |  | This study |
| yAF412 | GPDp-Hsp104-Pil1 + pRS416::Htt25Q | *MATa his3∆1 leu2∆0 ura3∆0 met15∆0::GPDp-HSP104-PIL1-CYC1t-KanMX pRS416::GPDp-Htt25Q-GFP(URA)* | yAF393 |  | This study |
| yAF413 | GPDp-Hsp104-Pil1 + pRS416::Htt103QP | *MATa his3∆1 leu2∆0 ura3∆0 met15∆0::GPDp-HSP104-PIL1-CYC1t-KanMX pRS416::GPDp-Htt103QP-GFP(URA)* | yAF393 |  | This study |
| yAF475 | GBP-Pea2 + Htt103QP-mCherry query strain for screen | *MATα can1Δ::STE2pr-LEU2 lyp1Δ his3Δ1 leu2Δ0 ura3Δ0 met15Δ::ADH1p-GBP-Pea2-KanMX pRS416::GPDp-Htt103QP-mCherry(URA)* | yAF470 |  | This study |
| yAF477 | tsa1Δ + ADH1p-Hsp104-GFP-Pea2 | *MATa tsa1Δ::KanMX4 his3Δ1 leu2Δ0 met15∆0::ADH1p-Hsp104-GFP-Pea2-CYC1t-NATMX ura3Δ0* | tsa1Δ | pAF070 | This study |
| yAF492 | Tsa1-GFP + GBP-Pea2 | MAT a ura3-52 leu2Δ1 trp1Δ63 his3Δ200 lys2ΔBgl hom3-10, ade2Δ1, ade8, hxt13Δ::URA3 TSA1(WT)_GFP::NAT/TRP1 met15Δ::ADH1p-GFPnanobody-Pea2-KanMX | SH164 | pAF118 | This study |
| yAF493 | Tsa1_C48S-GFP GBP-Pea2 | MAT a ura3-52 leu2Δ1 trp1Δ63 his3Δ200 lys2ΔBgl hom3-10, ade2Δ1, ade8, hxt13Δ::URA3 tsa1C48S-GFP::NAT/TRP1 cyh2 met15Δ::ADH1p-GFPnanobody-Pea2-KanMX | SH165 | pAF118 | This study |
| yAF494 | Tsa1_C171S-GFP GBP-Pea2 | MAT a ura3-52 leu2Δ1 trp1Δ63 his3Δ200 lys2ΔBgl hom3-10, ade2Δ1, ade8, hxt13Δ::URA3 tsaC171S-GFP::NAT/TRP1 cyh2 met15Δ::ADH1p-GFPnanobody-Pea2-KanMX | SH166 | pAF118 | This study |
| yAF495 | Tsa1_C48,171S-GFP GBP-Pea2 | MAT a ura3-52 leu2Δ1 trp1Δ63 his3Δ200 lys2ΔBgl hom3-10, ade2Δ1, ade8, hxt13Δ::URA3 tsa1C48S,C171S-GFP::NAT/TRP1 cyh2 met15Δ::ADH1p-GFPnanobody-Pea2-KanMX | SH169 | pAF118 | This study |
| yAF496 | Tsa1-GFP + GBP-Pea2 | MATα, his3Δ1, leu20, lysΔ0, ura3Δ0 Tsa1-GFP::hph met15Δ::ADH1p-GBP-Pea2-KanMX | SH192 | pAF118 | This study |
| yAF497 | Tsa1_DYF-GFP GBP-Pea2 | MATα, his3Δ1, leu20, lysΔ0, ura3Δ0 Tsa1DYF-GFP::hph met15Δ::ADH1p-GBP-Pea2-KanMX | SH193 | pAF118 | This study |
| yAF500 | ADH1p-Tsa1(S78D)-GFP-Pea2 | *MATa his3Δ1 leu2Δ0 met15Δ0::ADH1p-Tsa1(S78D)-GFP-Pea2-CYC1t-KanMX ura3Δ0* | BY4741 | pAF130 | This study |
| yAF501 | ADH1p-Hsp42(ΔCTD)-GFP-Pea2 | *MATa his3Δ1 leu2Δ0 met15Δ0::ADH1p-Hsp42(ΔCTD)-GFP-Pea2-CYC1t-KanMX ura3Δ0* | BY4741 | pAF131 | This study |
| yAF504 | ADH1p-Tsa1ΔOI1-GFP-Pea2 | *MATa his3Δ1 leu2Δ0 met15Δ0::ADH1p-Tsa1_ΔOI1-GFP-Pea2-CYC1t-KanMX ura3Δ0* | BY4741 | pAF136 | This study |
| yAF505 | ADH1p-Tsa1ΔOI2-GFP-Pea2 | *MATa his3Δ1 leu2Δ0 met15Δ0::ADH1p-Tsa1_ΔOI2-GFP-Pea2-CYC1t-KanMX ura3Δ0* | BY4741 | pAF137 | This study |
| yAF506 | ADH1p-Hsp42(ΔACD)-GFP-Pea2 | *MATa his3Δ1 leu2Δ0 met15Δ0::ADH1p-Hsp42(ΔACD)-GFP-Pea2-CYC1t-KanMX ura3Δ0* | BY4741 | pAF139 | This study |
| yAF507 | Tet1p-Hsp104GFP-Pea2 | URA3::CMV-tTA MATa his3Δ1 leu2Δ0 met15∆0::TET1p-Hsp104-GFP-Pea2-CYC1t-KanMX | R1158 | pAF133 | This study |
| yAF508 | Tet1p-GFP-Pea2 | URA3::CMV-tTA MATa his3Δ1 leu2Δ0 met15∆0::TET1p-GFP-Pea2-CYC1t-KanMX | R1158 | pAF142 | This study |
| yAF514 | ADH1p-Hsp42(ΔACD, ΔCTD)-GFP-Pea2 | *MATa his3Δ1 leu2Δ0 met15Δ0::ADH1p-Hsp42(ΔACD, ΔCTD)-GFP-Pea2-CYC1t-KanMX ura3Δ0* | BY4741 | pAF145 | This study |
| yAF517 | BY4741 + Hsp42(ΔNTD) | *MATa his3Δ1 leu2Δ0 met15Δ0::ADH1p-Hsp42(ΔNTD)-GFP-Pea2-CYC1t-KanMX ura3Δ0* | BY4741 | pAF149 | This study |
| yAF556 | BY4741 + Hsp42_CTD | *MATa his3Δ1 leu2Δ0 met15Δ0::ADH1p-Hsp42(CTD)-GFP-Pea2-CYC1t-KanMX ura3Δ0* | BY4741 | pAF154 | This study |
| yAF557 | BY4741 + Hsp42_ACD | *MATa his3Δ1 leu2Δ0 met15Δ0::ADH1p-Hsp42(ACD)-GFP-Pea2-CYC1t-KanMX ura3Δ0* | BY4741 | pAF155 | This study |
| yAF566 | Tet1p-Hsp104-GFP | URA3::CMV-tTA MATa his3Δ1 leu2Δ0 met15∆0::TET1p-Hsp104-GFP-Pea2-CYC1t-KanMX | R1158 | pAF171 | This study |
| yAF569 | Snf7-GFP + pYES2-Gal1p-Htt103QP-mCh | MATa his3∆1 leu2∆0 met15∆0 ura3∆0 Snf7-GFP-HIS3 pYES2-GAL1p-Htt103QP-mCherry(URA) | Snf7-GFP |  | This study |
| yAF570 | Pil1-GFP + pYES2-Gal1p-Htt103QP-mCh | MATa his3∆1 leu2∆0 met15∆0 ura3∆0 Pil1-GFP-HIS3 pYES2-GAL1p-Htt103QP-mCherry(URA) | Pil1-GFP |  | This study |
| yAJ002 | ADH1p-Hsp104-Pea2 + pYES2-Q25-EGFP | *MATa his3∆1 leu2∆0 ura3∆0 met15∆0::ADH1p-Hsp104-Pea2-CYC1t-KanMX pYES2-Q25-EGFP(URA)* | yAF275 |  | This study |
| yAJ003 | ADH1p-Hsp104-Pea2 + pYES2-QP25-EGFP | *MATa his3∆1 leu2∆0 ura3∆0 met15∆0::ADH1p-Hsp104-Pea2-CYC1t-KanMX pYES2-Qp25-EGFP(URA)* | yAF275 |  | This study |
| yAJ004 | ADH1p-Hsp104-Pea2 + pYES2-Q103-EGFP | *MATa his3∆1 leu2∆0 ura3∆0 met15∆0::ADH1p-Hsp104-Pea2-CYC1t-KanMX pYES2-Q103-EGFP(URA)* | yAF275 |  | This study |
| yAJ005 | ADH1p-Hsp104-Pea2 + pYES2-QP103-EGFP | *MATa his3∆1 leu2∆0 ura3∆0 met15∆0::ADH1p-Hsp104-Pea2-CYC1t-KanMX pYES2-Qp103-EGFP(URA)* | yAF275 |  | This study |
| yAJ008 | met15∆ vector control + pYES2-EGFP | *MATa his3∆1 leu2∆0 ura3∆0 met15∆0::KanMX pYES2-EGFP(URA)* | yAF290 |  | This study |
| yAJ009 | met15∆ vector control + pYES2-Q25-EGFP | *MATa his3∆1 leu2∆0 ura3∆0 met15∆0::KanMX pYES2-Q25-EGFP(URA)* | yAF290 |  | This study |
| yAJ010 | met15∆ vector control + pYES2-QP25-EGFP | *MATa his3∆1 leu2∆0 ura3∆0 met15∆0::KanMX pYES2-Qp25-EGFP(URA)* | yAF290 |  | This study |
| yAJ011 | met15∆ vector control + pYES2-Q103-EGFP | *MATa his3∆1 leu2∆0 ura3∆0 met15∆0::KanMX pYES2-Q103-EGFP(URA)* | yAF290 |  | This study |
| yAJ012 | met15∆ vector control + pYES2-QP103-EGFP | *MATa his3∆1 leu2∆0 ura3∆0 met15∆0::KanMX pYES2-QP103-EGFP(URA)* | yAF290 |  | This study |
| yAJ028 | GPDp-Hsp104-Pil1 (No GFP) + pYES2-EGFP | *MATa his3∆1 leu2∆0 ura3∆0 met15∆0::KanMX pRS416::GPDp-Htt103Q-GFP(URA) + pYES2-EGFP* | yAF393 |  | This study |
| yAJ029 | GPDp-Hsp104-Pil1 (No GFP) + pYES2-Q25-EGFP | *MATa his3∆1 leu2∆0 ura3∆0 met15∆0::KanMX pRS416::GPDp-Htt103Q-GFP(URA) + pYES2-Q25-EGFP* | yAF393 |  | This study |
| yAJ030 | GPDp-Hsp104-Pil1 (No GFP) + pYES2-QP25-EGFP | *MATa his3∆1 leu2∆0 ura3∆0 met15∆0::KanMX pRS416::GPDp-Htt103Q-GFP(URA) + pYES2-QP25-EGFP* | yAF393 |  | This study |
| yAJ031 | GPDp-Hsp104-Pil1 (No GFP) + pYES2-Q103-EGFP | *MATa his3∆1 leu2∆0 ura3∆0 met15∆0::KanMX pRS416::GPDp-Htt103Q-GFP(URA) + pYES2-Q103-EGFP* | yAF393 |  | This study |
| yAJ032 | GPDp-Hsp104-Pil1 (No GFP) + pYES2-QP103-EGFP | *MATa his3∆1 leu2∆0 ura3∆0 met15∆0::KanMX pRS416::GPDp-Htt103Q-GFP(URA) + pYES2-QP103-EGFP* | yAF393 |  | This study |
| Ydj1-GFP-FS | Ydj1-GFP-FS | MATa his3∆1 leu2∆0 met15∆0 ura3∆0 YDJ1-GFP-FS-HIS3 |  |  | Moreno et al. (2019) |
